# Rapid expansion of SARS-CoV-2 variants of concern is a result of adaptive epistasis

**DOI:** 10.1101/2021.08.03.454981

**Authors:** Michael R. Garvin, Erica T. Prates, Jonathon Romero, Ashley Cliff, Joao Gabriel Felipe Machado Gazolla, Monica Pickholz, Mirko Pavicic, Daniel Jacobson

**Affiliations:** Oak Ridge National Laboratory, Computational Systems Biology, Biosciences, Oak Ridge, TN; National Virtual Biotechnology Laboratory, US Department of Energy; The Bredesen Center for Interdisciplinary Research and Graduate Education, University of Tennessee Knoxville, Knoxville, TN; Departamento de Física, Facultad de Ciencias Exactas y Naturales, Universidad de Buenos Aires, Buenos Aires, Argentina; Instituto de Física de Buenos Aires (IFIBA), CONICET-Universidad de Buenos Aires, Buenos Aires, Argentina

**Keywords:** SARS-CoV-2, recombination, haplotype, epistasis, variants of concern, syncytia

## Abstract

The SARS-CoV-2 pandemic recently entered an alarming new phase with the emergence of the variants of concern (VOC) and understanding their biology is paramount to predicting future ones. Current efforts mainly focus on mutations in the spike glycoprotein (S), but changes in other regions of the viral proteome are likely key. We analyzed more than 900,000 SARS-CoV-2 genomes with a computational systems biology approach including a haplotype network and protein structural analyses to reveal lineage-defining mutations and their critical functional attributes. Our results indicate that increased transmission is promoted by epistasis, i.e., combinations of mutations in S and other viral proteins. Mutations in the non-S proteins involve immune-antagonism and replication performance, suggesting convergent evolution. Furthermore, adaptive mutations appear in geographically disparate locations, suggesting that either independent, repeat mutation events or recombination among different strains are generating VOC. We demonstrate that recombination is a stronger hypothesis, and may be accelerating the emergence of VOC by bringing together cooperative mutations. This emphasizes the importance of a global response to stop the COVID-19 pandemic.

## 1. Introduction

In late 2020, three SARS-CoV-2 variants of concern (VOC); B.1.1.7 (Alpha), B.1.351 (Beta), and P.1 (Gamma), rapidly spread due to enhanced transmission rates and have since been linked to increased hospitalizations and mortalities [1–9]. In early 2021, several new VOC appeared including, B.1.617 (Delta), B.1.427 (Epsilon), and B.1.526 (Iota). B.1.617 initiated the COVID-19 crisis in India [10] and is now causing the majority of new infections worldwide, even in geographic regions with robust sampling and early detection. There is therefore a critical need to identify accurate predictors and biological causes for the emergence of VOC to predict future ones.

Towards that end, two major weaknesses of recent efforts need to be addressed. First, the predominant mutations used to identify and describe the VOC are in the spike glycoprotein (S) whereas those in other genomic regions are largely ignored, but they are almost certainly biologically important. Second, species-level tools are predominantly used to study the molecular evolution of SARS-CoV-2, but existing population-level tools are a better fit for addressing the important questions related to the VOC [11–13] (Box 1). This has significant implications for societal responses to the ongoing pandemic. For example, the D614G and N501Y substitutions in S, associated with increased virulence, have appeared in geographically distant regions, but their emergence patterns indicate that the mechanisms of their evolution significantly differ. Such observations are not straightforward using species-level approaches. For D614G, this pattern appears to be a result of mutation followed by simple inheritance, but the case of N501Y is less clear [14] (Figure 1). Its appearance in different VOC could be the result of independent, repeated mutational events, or recombination, which is a common mechanism to accelerate evolution in positive strand RNA viruses such as SARS-CoV-2. This process can occur from co-infection of multiple viral strains in a single individual and has recently been demonstrated in North America and the United Kingdom (U.K.) [15–19]. If the VOC were generated by recombination then the movement of individuals from different geographic regions may be rapidly accelerating the evolution of the virus [17, 20].

**Fig. 1.**
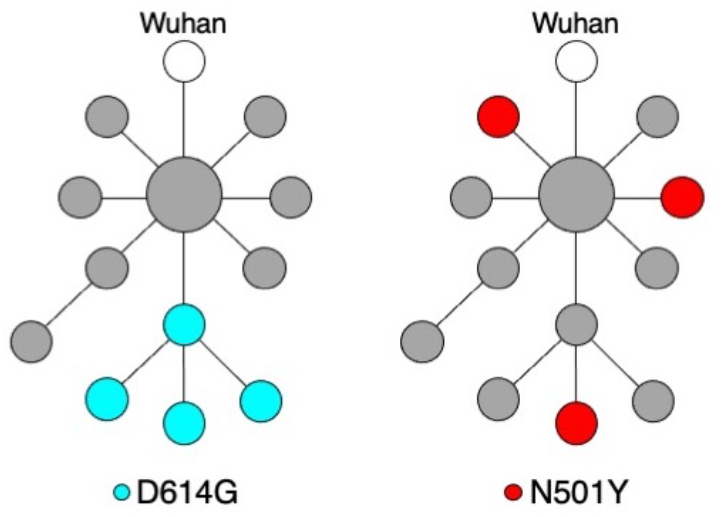
Haplotype networks representing different genealogies of mutations. Each node represents a haploid sequence (haplotype) and the edge between each node indicates a mutational event leading to a new haplotype much like a family pedigree. Node sizes reflect population frequencies. Mutations can be passed down by descent after a single mutational event as was the case for the D614G mutation in the spike glycoprotein (S) (left). Conversely, N501Y in S appears in different genealogical lineages, which could be from independent mutation events, or from recombination among different viral strains co-infecting an individual (right). Determining which process is occurring has significant implications for addressing current variants of concern and predicting future ones.

Here, we processed more than 900,000 SARS-CoV-2 genomes and applied a median-joining-network (MJN) algorithm to generate a network derived from linked sets of mutations (a haplotype) and combined those results with protein structural analyses [21, 22]. The haplotype network indicates VOC arose and spread rapidly as a result of specific combinations of mutations (epistasis) in S and non-S proteins. Critically, these VOC spread in a manner consistent with either repeat mutations at adaptive sites or recombination events rather than simple inheritance. We explore the individual effects of key mutations in the SARS-CoV-2 proteome that are shared among different VOC. Specifically, we identify a signature of co-evolution and the molecular basis for it between residue 501 in S and the host ACE2. A test of the hypothesis that repeat mutations are responsible for late 2020 VOC indicates recombination may be a more likely mechanism for their generation and warrants further scrutiny. This work emphasizes the importance of determining the mechanisms of community spread in generating future VOC [23].

## 2. Results and Discussion

### 2.1. Identification of lineage-defining mutations of VOC

We processed more than 900,000 human and mink SARS-CoV-2 genomes, built a haplotype network using the 640,211 that passed QC, and annotated with PANGO lineages from GISAID (Figure 2 and Supplementary File). This genealogy-based approach to molecular evolution identifies the mutations that define VOC and variants of interest (VOI) based on the edge that initiates their corresponding clusters of nodes (Table 1).

**Fig. 2.**
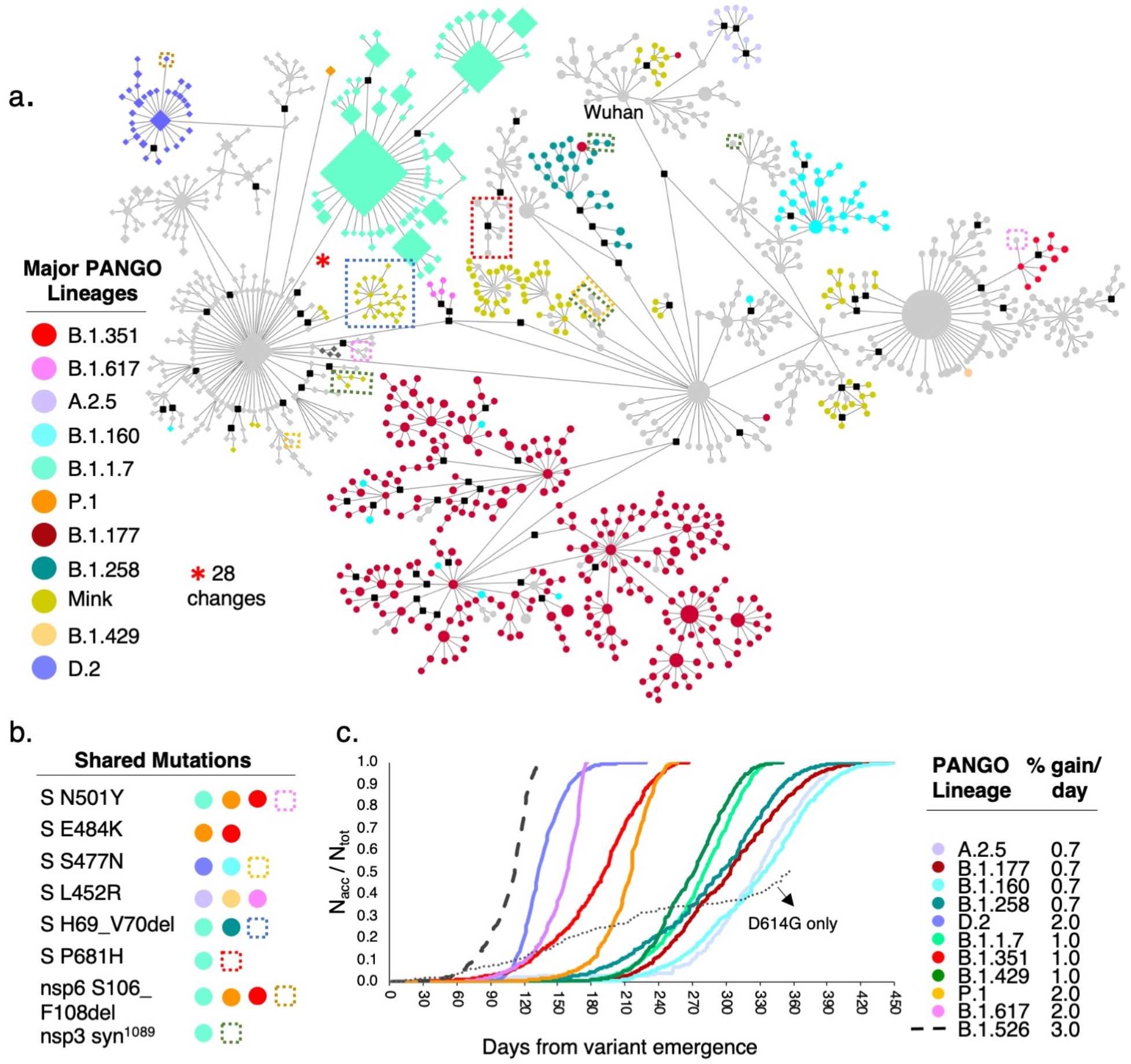
Haplotype network of SARS-CoV-2 genomes. **a.** MJN-based network built from haplotypes found in more than 25 individuals (N=640,211 sequences) using 2,128 variable sites. Nodes indicate a haplotype and size reflects frequency. Edges denote ordered mutation events that may be a single change or many at one time. Colors identify PANGO lineages from GISAID. Diamond-shaped nodes correspond to haplotypes carrying a three nucleotide deletion in the nucleocapsid gene (N) at sites 28881-28883 (R203K and G203R). Black square nodes are inferred haplotypes, dashed-line boxes define subgroups of haplotypes within a lineage with a disjoint mutation that is also found in VOC that could indicate repeat mutation events or recombination. Red asterisk indicates the edge leading to B.1.1.7 and represents 28 nucleotide changes discussed in text. Several lineages show introgression from others (e.g., cyan nodes, B.1.160, into brick red, B.1.177). We note that P.1 and B.1.351 are under-sampled compared to B.1.1.7. **b.** Several important mutations in S and non-S proteins appear in multiple variants of concern (VOC). For example, the B.1.1.7 variant carries four mutations that are in disjoint haplotypes: S N501Y, S P681H, a silent mutation in the codon for amino acid 1089 in nsp3, and the S H69_V70del that is also found in a clade of haplotypes from mink, identified by the blue dashed-line box in (a). **c.** Accumulation rate for common GISAID lineages including VOC and likely (L-VOC) represented by the ratio between the accumulated number of reported sequences of a given lineage per day since the appearance of that haplotype (N_acc_) divided by the corresponding total number (N_tot_) at the final sample date for this study (April 2021). Colors of curves correspond to node colors in (a). All VOC display accumulation rates of at least 1% of the total for that variant per day. The remaining are less than 1%, except for the VOI B.1.526 (not displayed in the network), with the highest rate of 3% per day, indicating that further scrutiny of this variant is warranted. We also plotted the accumulation rate for lineages that carry the widely reported S D614G mutation but without the nsp12 P323L commonly found with it, supporting our previous hypothesis [21] that mutations in S alone are not responsible for the rapid transmission of these VOC/VOI but is resulting from epistasis among S and non-S mutations. Reference sequence: NC_045512, Wuhan, December 24, 2019.

**Table 1.**
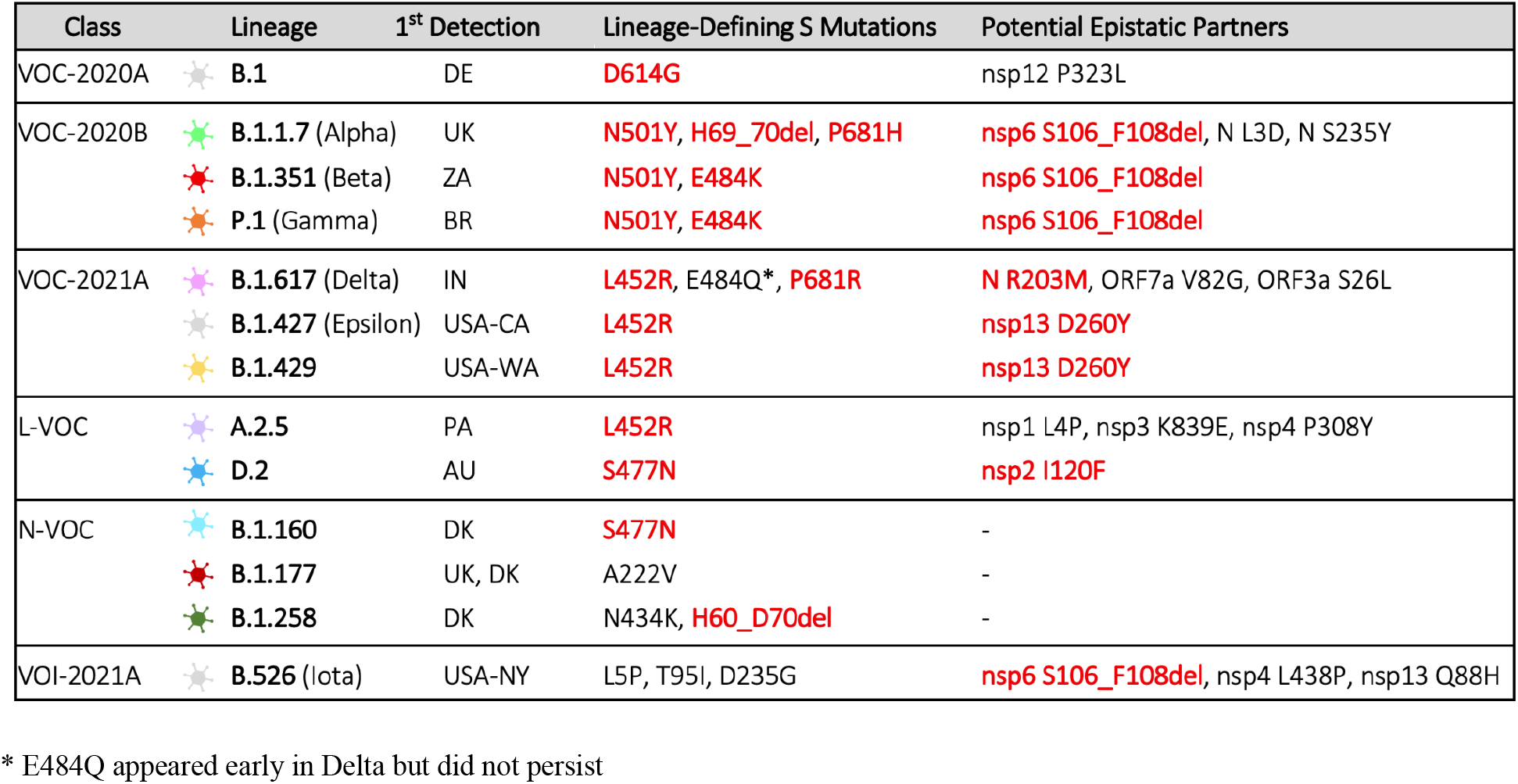
Major lineages shown in the haplotype network and their defining mutations. Potential epistatic mutations in S and non-S proteins that define SARS-CoV-2 lineages discussed here are shown. VOC/VOI are grouped in classes according to their peak time of emergence using the notation VOC/VOI-(year and period of emergence), with A and B corresponding to the first and last six months of the year, respectively. The nomenclature of the lineages used by the Center for Disease Control (CDC) are listed. L-VOC denotes likely variants of concern, that is, those that we propose to have strong potential to become VOC. Non-VOC (N-VOC) are not identified by CDC as VOC. The functional impact of mutations highlighted in red are discussed in the text.

| Class | Lineage | 1<sup>st</sup> Detection | Lineage-Defining S Mutations | Potential Epistatic Partners |
| --- | --- | --- | --- | --- |
| VOC-2020A | B.1 | DE | D614G | nsp12 P323L |
| VOC-2020B | B.1.1.7 (Alpha) | UK | N501Y, H69_70del, P681H | nsp6 S106_F108del, N L3D, N S235Y |
|  | B.1.351 (Beta) | ZA | N501Y, E484K | nsp6 S106_F108del |
|  | P.1 (Gamma) | BR | N501Y, E484K | nsp6 S106_F108del |
| VOC-2021A | B.1.617 (Delta) | IN | L452R, E484Q*, P681R | N R203M, ORF7a V82G, ORF3a S26L |
|  | B.1.427 (Epsilon) | USA-CA | L452R | nsp13 D260Y |
|  | B.1.429 | USA-WA | L452R | nsp13 D260Y |
| L-VOC | A.2.5 | PA | L452R | nsp1 L4P, nsp3 K839E, nsp4 P308Y |
|  | D.2 | AU | S477N | nsp2 I120F |
| N-VOC | B.1.160 | DK | S477N | - |
|  | B.1.177 | UK, DK | A222V | - |
|  | B.1.258 | DK | N434K, H60_D70del | - |
| VOI-2021A | B.526 (Iota) | USA-NY | L5P, T95I, D235G | nsp6 S106_F108del, nsp4 L438P, nsp13 Q88H |

\* E484Q appeared early in Delta but did not persist

This approach also facilitates the identification of features common to different VOC/VOI, which can reveal molecular events underlying their rapid spread. For example, the VOC reported in late 2020 (VOC-2020B), B.1.1.7, B.1.351, and P.1, can be defined by a triple amino acid deletion in the nonstructural protein 6 (nsp6; S106_F108del) and the substitution N501Y in S. Notably, in contrast to the latter, very little attention has been dedicated to unravel the biological importance of the mutation in nsp6 (Figure 2a, Table 1). In addition, because the MJN algorithm produces time-ordered mutation series and the frequency of specific haplotypes, our analysis reveals a likely VOC (L-VOC, Table 1) that appeared in April 2020, but was missed. The D.2 variant in Australia was defined by an initial I120F substitution in nsp2 followed by S477N in S, which led to its rapid expansion. In contrast, B.1.160, which carries S S477N solely, did not rapidly expand. This underscores the usefulness of the haplotype network, as it is able to simultaneously convey the timing of mutational events and the frequency of the resulting linked set of mutations, i.e., haplotypes.

A disproportionate number of all haplotypes, including B.1.1.7 and D.2, are present in GISAID from the UK and Australia. In order to account for this sampling bias), we plotted the cumulative number of different variants and compared the resulting slopes of the linear range of the curves (Figure 2c). VOC-2020B and VOC/VOI reported in early 2021 (VOC-2021A and VOI-2021A, respectively; Table 1) display higher daily accumulation rates compared to other variants, e.g. B.1.177, which show less than 1% accumulation per day. Notably, the rapid increase in D.2 (2%), but not B.1.160 (0.7%), supports our interpretation of the haplotype network defining it as a L-VOC and the major role of epistasis (here S S477N and nsp2 I120F). These analyses reinforce the importance of monitoring these variants closely and of identifying haplotypes based on both S and non-S mutations .

### 2.2. Recombination may accelerate the emergence of SARS-CoV-2 VOC

Haploid, clonally replicating organisms such as SARS-CoV-2, are predicted to become extinct due to the accumulation of slightly deleterious mutations, i.e., Muller’s ratchet [24]. Recombination is a rescue from Muller’s ratchet and can accelerate evolution by allowing for the union of advantageous mutations from divergent haplotypes [15, 25]. In SARS-CoV-2, recombination manifests as a template switch during replication when more than one haplotype is present in the host cell, i.e., the virus replisome stops processing a first RNA strand and switches to a second one from a different haplotype, producing a hybrid [16]. In fact, because template switching is a necessary step during the negative-strand synthesis of SARS-CoV-2 [26] and is a major mechanism of coronavirus evolution [27], it would be improbable for recombination *not* to occur in the case of multiple strains infecting a cell [28], especially given that more than 1 billion copies of the virus are generated in an individual during infection [29].

Several VOC-2020B exhibit large numbers of new mutations relative to any closely related sequence indicating rapid evolution of SARS-CoV-2 (Figure 3a) [30]. For example, the first emergent haplotype of B.1.1.7 differs from the most closely related haplotype by 28 nucleotide changes (red asterisk, Figure 2a). Notably, all 28 differences between the Wuhan reference and B.1.1.7 appeared in earlier haplotypes and, in some cases, in multiple lineages (Table S1) indicating that either repeat mutations at a site or recombination were instrumental to the emergence of VOC-2020B.

**Fig. 3.**
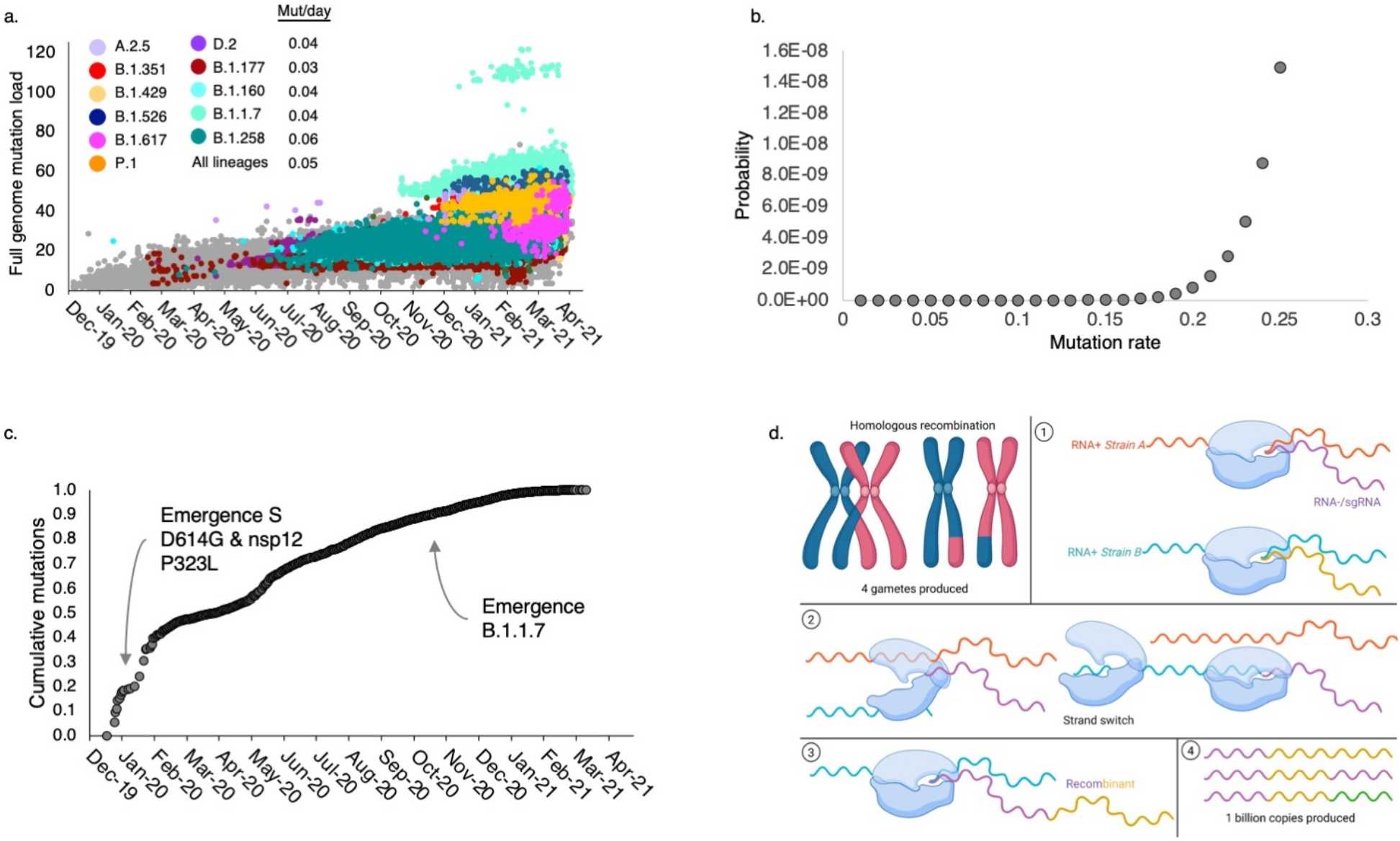
Mutation rates and genome-wide mutation load of SARS-CoV-2. **a.** A rapid increase in the number of mutations per individual genome is evident in the VOC-2020B (B.1.1.7, B.1.351, and P.1). The outliers of the B.1.1.7 lineages (mint green) are a subset of that lineage due to a single, 57 nucleotide deletion in ORF7a (amino acids 5-23). Mutation rate based on a linear regression of genome-wide mutation load for five common lineages and all lineages varies between 0.04 and 0.06 mutations per day. **b.** Probability estimate for 13 mutations occurring in a single event as a function of mutation rate. **c.** Population-level analysis of new mutations per day (accounting for multiple mutations per site) displays a declining rate of mutations after the emergence of B.1 (S D614G and nsp12 P323L). A slight increase around the emergence of D.2 in Australia (May 2020) is also evident. There is not an increase in rate with the emergence of B.1.1.7 that could explain the rapid accumulation of mutations shown in (a). **d**. Contrast between recombination in a diploid organism (four haploid gametes) and in RNA+ *betacoronaviruses*. In a diploid organism, recombinants are produced from the crossover of homologous chromosomes during meiosis. In SARS-CoV-2, recombination occurs when two or more strains (haplotypes) infect a single cell (1). The replisome dissociates (2) from one strand and switches to another, (3) generating a hybrid RNA. The resulting chimera (4) can be as simple as a section of *strain A* (pink) fused to a section of *strain B* (yellow) or more complex recombinants if strand switching occurs more than once or there are multiple strains per cell (green). 1 billion copies are estimated in an individual during infection [27].

We tested the hypothesis that these numerous differences arose from repeat, independent mutations. The majority of the 28 differences (15 of the 28) are deletions that could be considered two single mutational events, as does a 3-bp change in N (28280-22883) since they occur in factors of three (a codon), maintaining the coding frame. The two deletions and the full codon 3-bp change in N combined with the 10 remaining single-site mutations results in the conservative estimate of 13 mutations. The plot of accumulating mutations reveals a linear growth of roughly 0.05 mutations per day based on all lineages (Figure 3a) and therefore it would take 260 days for 13 mutational events to occur. We note that the mutation rate of 0.05 per day is based on neutral, slightly deleterious (common in expanding populations), and adaptive sites of the entire viral population and is therefore a robust unbiased estimate. Furthermore, it is conservative because in many cases the B.1.1.7-defining sites appear to have mutated not twice, but several times (Table S1), and always to the same nucleotide state. Considering this hypothesis, the appearance of B.1.1.7 in October 2020 would require it to have emerged in January 2020 and yet the nearest haplotype harboring S S501Y was not sampled until June 2020 and no intermediate haplotypes have been identified. Likewise, the probability that the 13 mutational events occurred between June and October is 1×10^-15^ (Supplemental Methods). Even with a mutation rate that is five-fold higher than our estimate (Figure 3a), the occurrence of 13 mutations in a single event is highly improbable (Figure 3c). Therefore, repeat mutation events do not explain the rapid increase in mutation load of B.1.1.7. Alternatively, the rapid accumulation of mutations in B.1.1.7 could be the result of an increased mutation rate just prior to its appearance. To test this hypothesis, we plotted the population-level mutations per day, including repeat mutations at variable sites, which did not reveal any increase in mutation rate at the time of B.1.1.7 emergence, but instead it displayed a decrease with its emergence (Figure 3c, Figure S1).

Another commonly proposed hypothesis is that the large increase in mutations could have occured in a few particular individuals with immunodeficiency disorders [30]. However, that would have to occur on multiple continents to explain B.1.1.7, B.1.351, and P.1 because these and other haplotypes also show rapid increases in mutation loads at this time (late 2020). In addition, a virus in an immunocompromised individual would be under no selective pressure and mutations would therefore be due to drift and not selection. However, another possibility is that adaptive mutations are generated from repeat infections in human or non-human hosts, and immunocompromised individuals are providing an environment highly conducive to infections from multiple strains and subsequent recombination.

Recombination among divergent haplotypes (Figure 3d) is the most parsimonious explanation for the rapid increase in mutation load in multiple VOC given (1) the absence of a substantial increase in mutation rate at any time prior to the appearance of the VOC along with their increased mutation load, which often occurs from template-switching during replication [16], (2) the widespread and early circulation of the majority of the mutations associated with them in other haplotypes and, (3) that several mutations appear in haplotypes that are clearly disjoint in the network (Figure 2a). For example, N501Y and P681H in S appear in several divergent haplotypes, including one mink subgroup from Denmark and a basal node to B.1.351 (without the nsp6 deletion found in VOC-2020B).

It has been argued that multiple independent mutation events in S (N501Y, S477N, and P681H) are the result of positive selection [14]. However, mutation is agnostic to selection, i.e., even if sites appearing in multiple lineages are under positive selection, their appearance in disjoint haplotypes still requires repeated mutations in the absence of recombination, which we demonstrated to be unlikely. In addition, B.1.1.7 also carries a mutation in nsp3 that appears in disjoint haplotypes (including mink) but is unlikely to be under selection because it is synonymous, weakening the argument that adaptive evolution and increased mutation rate at a site are linked (although synonymous mutations could be adaptive in some cases).

In addition, recombination can generate a high number of false-positive tests of positive selection [31], and the complexity of coronavirus recombinants compared to those generated in diploid organisms through homologous chromosome crossovers (Figure 3d) makes that process difficult to detect. Therefore, tests for positive selection based on multiple independent mutations at a site may, in fact, be false positives due to recombination [14]. Alternatively, the majority of the B.1.1.7 mutations could be explained by the admixture and recombination among lineages and in support of this, a random scan of 100 FASTQ files from B.1.1.7 available in the NCBI SRA database identified two co-infected individuals (Table S2).

The presence of recombinants of SARS-CoV-2 was demonstrated in a recent analysis of sequences in the U.K. that leveraged the geographical and temporal presence of specific sequences whose parents appeared to be from B.1.1.7 and B.1.177-derived strains [19]. Although this demonstrates that recombination is occurring, it does not answer if B.1.1.7 itself arose through recombination. As noted above, the majority of B.1.1.7-defining mutations appeared earlier in the pandemic outside the U.K. (Table S1) and therefore approaches that focus on a single geographic location will miss parental variants if they came from elsewhere. Identification of recombinants using the method of Jackson et. al. [19] is even more difficult if the parental lineages were less fit or rare. The haplotype network here suggests that this is likely the case, given the low frequency of haplotypes that carry a VOC-defining mutation in S compared to haplotypes that gain an additional non-S mutation that displays increased frequency due to epistasis and thus could be sampled . Large-scale analyses with newer methods that include rare variants and a global distribution may be able to determine if B.1.1.7 and other VOC-2020B arose through recombination that resulted in this higher mutation load [32].

### 2.3. Key mutations in S target enhanced cellular invasion and neutralizing antibody escape

Given our results and previous hypothesis [21] that epistasis is responsible for increased transmission of VOC [37], we performed protein structural analyses and discussed the functional effects of likely key mutations in S and non-S proteins shared by several VOC (Table 1). Overall, VOC-shared mutations in S are mostly associated with an improved capacity of entering host cells and of escaping neutralizing antibodies (Table 2).

**Table 2.**
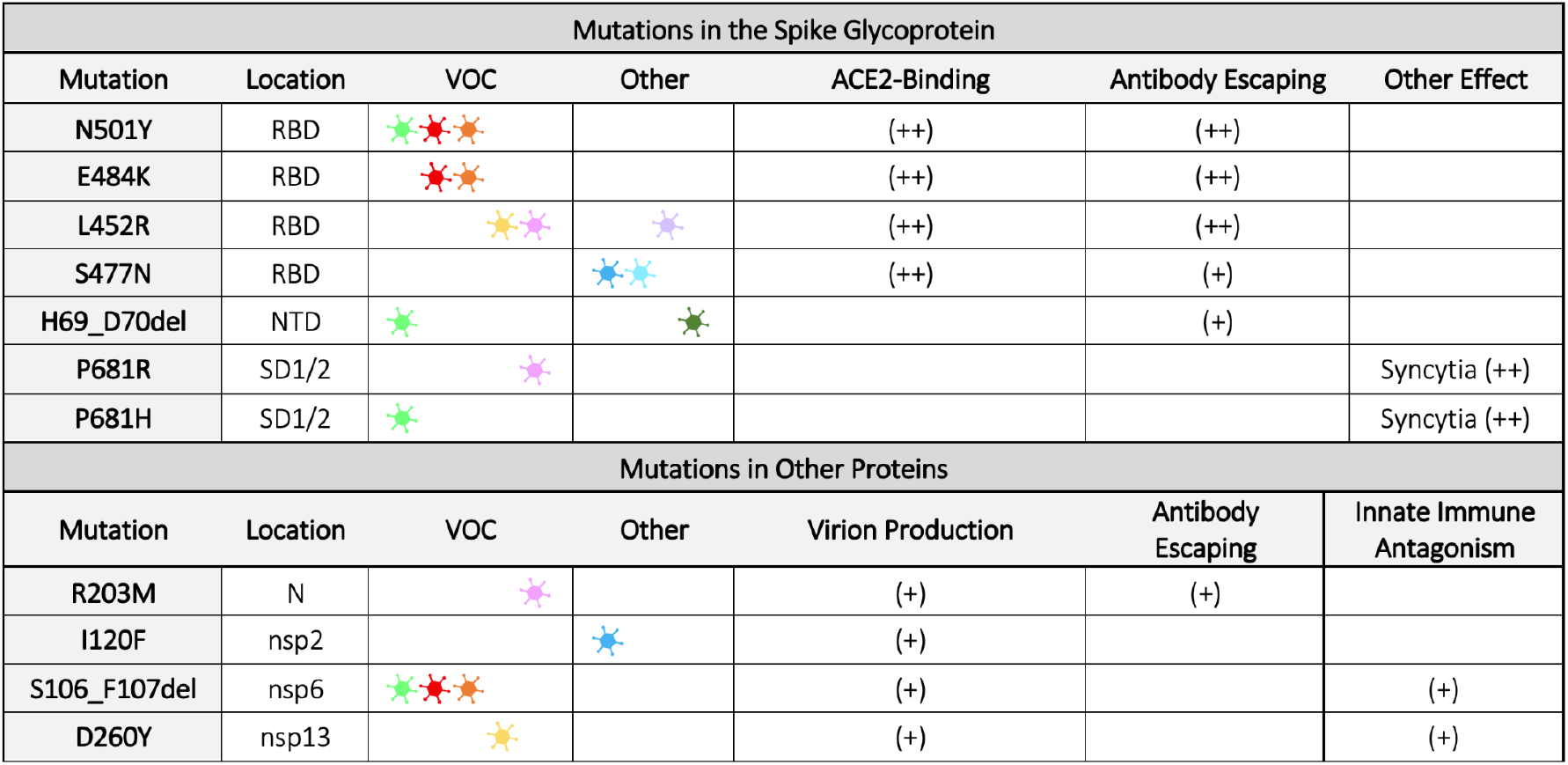
Summary of the predicted or determined functional impact of VOC-shared mutations. Likely key VOC-shared mutations, highlighted in Table 1, are analyzed. SARS-CoV-2 lineages are represented as symbols using the color code introduced in Figure 1. Mutations in S are associated with enhanced binding-affinity to host ACE2, neutralizing antibody escape, and increased syncytia formation. Mutations in non-S proteins might change virion production and enhance antagonism to host innate immune responses. A confidence label is assigned to these functional effects as (+) predicted from structural analysis or (++) inferred from experimental evidence. In S, the receptor-binding domain (RBD), N-terminal domain (NTD), and subdomains 1 and 2 (SD1/2) are specified.

| Mutations in the Spike Glycoprotein |  |  |  |  |  |  |
| --- | --- | --- | --- | --- | --- | --- |
| Mutation | Location | VOC | Other | ACE2-Binding | Antibody Escaping | Other Effect |
| N501Y | RBD |  |  | (++) | (++) |  |
| E484K | RBD |  |  | (++) | (++) |  |
| L452R | RBD |  |  | (++) | (++) |  |
| S477N | RBD |  |  | (++) | (+) |  |
| H69_D70del | NTD |  |  |  | (+) |  |
| P681R | SD1/2 |  |  |  |  | Syncytia (++) |
| P681H | SD1/2 |  |  |  |  | Syncytia (++) |
| Mutations in Other Proteins |  |  |  |  |  |  |
| Mutation | Location | VOC | Other | Virion Production | Antibody Escaping | Innate Immune Antagonism |
| R203M | N |  |  | (+) | (+) |  |
| I120F | nsp2 |  |  | (+) |  |  |
| S106_F107del | nsp6 |  |  | (+) |  | (+) |
| D260Y | nsp13 |  |  | (+) |  | (+) |

**S D501Y** -Site 501 located in the receptor-binding domain (RBD) of SARS-CoV-2 S plays a major role in the affinity of the virus to the host receptor, ACE2 [38]. Structural analyses highlighted the importance of the interactions with human ACE2 (hACE2) near the site 501 of the RBD of S, particularly via a sustained H-bond between RBD N498 and hACE2 K353 [39]. Naturally occurring mutations at site 501, N501Y and N501T are reported to increase affinity to hACE2 [40] and this site is located near a linear B cell immunodominant site [35]. Therefore, the mutation may allow SARS-CoV-2 variants to escape neutralizing antibodies (Figure 4, Figure 5), which is supported by reports of reduced activity against pseudotyped viruses carrying this mutation [41].

**Fig. 4.**
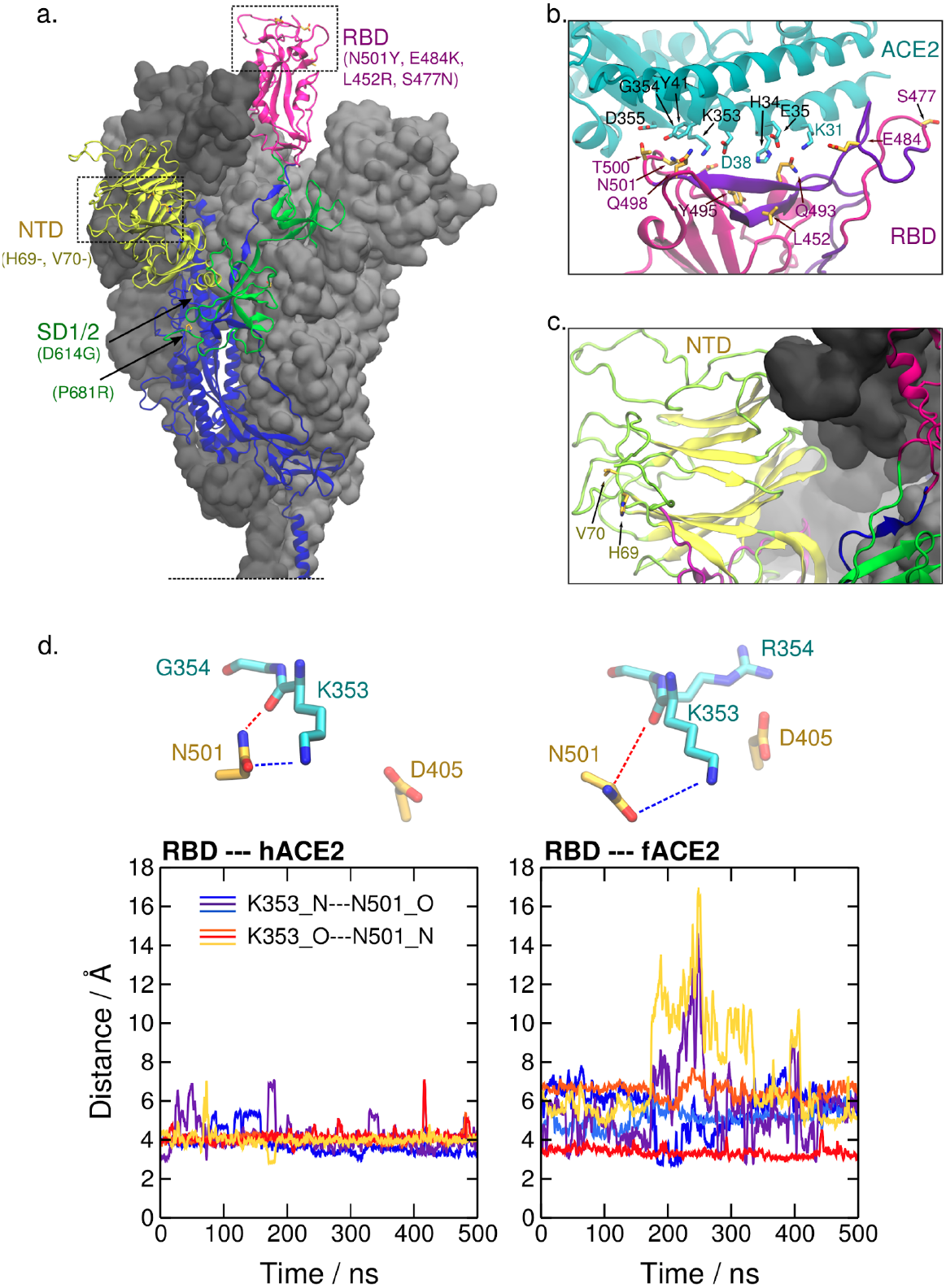
Location of mutation sites of SARS-CoV-2 VOC on the structure of the spike glycoprotein. **a.** Several mutations associated with dominant haplotypes are located in the receptor-binding domain (RBD, aa. 331-506), N-terminal domain (NTD, aa. 13-305), and subdomains 1 and 2 (SD1/2, aa. 528-685) of S. The structure of S in the prefusion conformation derived from PDB ID 6VSB [33] and completed *in silico* [34] is shown. Glycosyl chains are not depicted and the S trimer is truncated at the connecting domain for visual clarity. The secondary structure framework of one protomer is represented and the neighboring protomers are shown as a gray surface. **b.** Mutation sites in the S RBD of SARS-CoV-2 VOC, such as 484, 452, 477, and 501 are located at or near the interface with ACE2. Notably, site 452 and 484 reside in an epitope that is a target of the adaptive immune response in humans (aa. 480-499, in violet) and site 501 is also located near it [35]. Dashed lines represent relevant polar interactions discussed here. PDB ID 6M17 was used [36]. **c.** The sites 69 and 70 on the NTD, which are deleted in the VOC B.1.1.7, are also found near an epitope (aa. 21-45, in violet) [35]. **d.** Time progression of N---O distances between atoms of N501 in RBD and K353 in human and ferret ACE2 (hACE2 and fACE2, respectively) from the last 500 ns of the simulation runs. Colors in the plots correspond to the distances K353_N---N501_O (cold colors) and K353_O---N501_N (warm colors) in three independent simulations of each system. These distances are represented in the upper part of the figure.

**Fig. 5.**
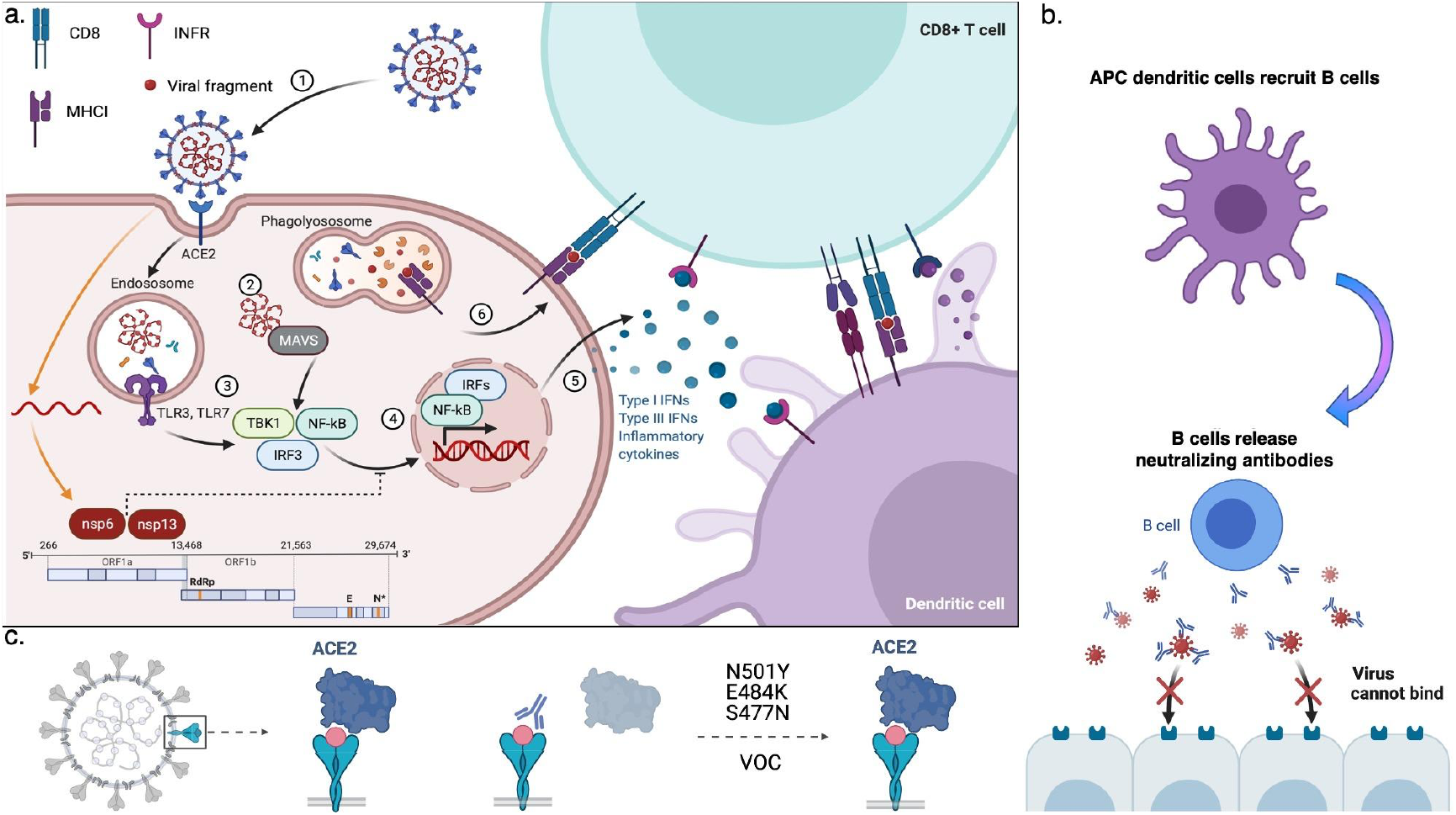
Host response to viral infection. **a**. As part of the innate immune response, (Step 1) the SARS-CoV-2 virus is internalized into endosomes and degraded. (Step 2) viral RNA activates the mitochondrial antiviral innate immunity (MAVS) pathway and (Step 3) degraded proteins activate the toll receptor pathway (TLR3/TLR7), which result in the (Step 4) phosphorylation of TBK1 and translocation of NF-*k*B and IRF3 to the nucleus, where they regulate the transcription of immune genes including interferons (IFNs, Step 5). IFNs recruit CD8+ T cells that, (Step 6) recognize fragments of the virus on the cell surface via their class I major histocompatibility complex (MHC I) receptors and are activated by dendritic cells (antigen processing cells, or APC). If the virus bypasses innate immunity (orange arrows) nonstructural proteins (nsp6 and nsp13) block the IRF3 nuclear translocation. **b.** APCs recruit B lymphocytes and stimulate the production of antibodies that recognize SARS-CoV-2 S (whereas T cells recognize fragments of S bound to MHC I). **c.** The neutralizing antibodies block binding of the virus to the ACE2 receptor and can prevent re-infection but mutations in the receptor-binding domain (RBD), e.g., S Asn^501^Tyr, prevent binding of the antibodies and the virus is then able to bind the receptor again even if individuals experienced exposure to an earlier strain or were vaccinated. *Created with BioRender.com*.

Cross-species transmission of SARS-CoV-2 provides further information on the selectivity of site 501. Repeated infection of mice with human SARS-CoV-2 selected for an S Y501 [42] that likely results in stabilization of the RBD-ACE2 interaction via π-stacking of Y501 with Y41 in ACE2 (Figure 4a-b). Several introductions into farmed mink (*Neovison vison*), which caused a substantial increase in their mortality [43], have not led to the same selection. To date, reported sequences in GISAID of SARS-CoV-2 in this host carry either S N501 (prevalent), or S T501, which appeared in independent farms (Table S3) [43]. In ACE2 of these taxa, Y41 is conserved but a differing, positively charged amino acid is found nearby (H354). The RBD 501-binding region of the ACE2 is highly conserved, except for site 354, suggesting its important role in viral fitness (Table 3). Ferrets and pangolins carry a large basic residue at site 354 (arginine and histidine, respectively). Absence of transmissions to ferrets even after long-term exposure suggests that ferret infection may require improved viral fitness [44]. In agreement with this, it was reported that the adaptive substitution N501T was detected in all infected ferrets in the laboratory [45].

**Table 3.** Surface exposed residues of ACE2 orthologues forming the region of contact with site 501 of SARS-CoV-2 S. Relative to the human sequence, almost all these residues are either conserved (“|”) or replaced by a nearly equivalent amino acid in mouse, American mink, European mink, ferret, and pangolin. Notably, there is a nonconservative substitution of G354 to a bulky positively charged amino acid in most species. Our structural analyses suggest that this substitution contributes to a putative host-dependent selective pressure at site 501 of SARS-CoV-2 S. Prevalent residues reported at this site are informed in order of frequency.

| Species | Residues in ACE2 |  |  |  |  |  |  |  |  | S 501 |
| --- | --- | --- | --- | --- | --- | --- | --- | --- | --- | --- |
| *Homo sapiens* (Human) | D38 | Y41 | Q42 | L45 | K353 | G354 | D355 | R357 | I358 | Y, N |
| *Mus musculus* (House mouse) |  |  |  |  | H |  |  |  |  | Y, N |
| *Neovison vison* (American mink) | E |  |  |  |  | H |  |  |  | N, T |
| *Mustela lutreola* (European mink) | E |  |  |  |  | R |  |  |  | N, T |
| *Mustela putorius furo* (Ferret) | E |  |  |  |  | R |  |  |  | T, N |
| *Manis pentadactyla* (Pangolin) | E |  |  |  |  | H |  |  |  | N, T |

To further investigate the role of the G354 versus R354 in the adaptive mutation of site 501 in S, we performed simulations of the truncated complexes of N501-carrying RBD of SARS-CoV-2 and ACE2, from human (hACE2) and ferret (fACE2). The simulations indicate remarkable differences between the two systems in the region surrounding site 501. We identified that the main ACE2 contacts with N501 were the same for both species, namely, Y41, K353, and D355, but the frequency of these contacts is lower in the simulations of fACE2 (Table S4).

We then analyzed structural features of ACE2 K353 and RBD N501 interactions. Atom distances computed from the simulations indicate a weaker electrostatic interaction between this pair of residues in ferret compared to human (Figure 4d). This effect is accompanied by a conformational change of fACE2 K353. In ferret, the side chain of K353 exhibits more stretched conformations, i.e., a higher population of the *trans* mode of the dihedral angle formed by the side chain carbon atoms (Figure S2). This conformational difference could be partially attributed to the electrostatic repulsion between the bulky positively charged amino acids, K353 and R354. Additionally, the simulations suggest a correlation, in a competitive manner, between other interactions that these residues display with the RBD. For example, the salt bridge fACE2_R354---RBD_D405 and the H-bond interaction fACE2_K353---RBD_Y495 alternate in the simulations (Figure S3). This also suggests that the salt bridge formed by fACE2 R354 drags K353 apart from RBD N501, weakening the interaction between this pair of residues in ferrets.

These analyses indicate that site 354 in ACE2 significantly influences the interactions with RBD in the region of site 501 and is likely playing a major role in the selectivity of its size and chemical properties. We propose that, in contrast to Y501, a smaller H-bond-interacting amino acid at site 501 of RBD, such as the threonine reported in minks and ferrets, may optimize interactions on the region, e.g., the salt bridge fACE2_R354---RBD_D405. The differences in the region of ACE2 in contact with site 501 seem to have a key role for host adaptation and further investigation may reveal details of the origin of COVID-19.

**S E484K** - A recent study suggested that E484 exhibits intermittent interactions with K31 in ACE2 [39]. This mutation is associated with higher affinity to ACE2 [40], which may be explained by its proximity to E75 in ACE2 and possibly the formation of a salt bridge. In addition, this site is part of a linear B cell immunodominant site [35] and S E484K was shown to impair antibody neutralization [41]. This mutation is typically associated with the B.1.351 and P.1 VOC, but it also appeared in some B.1.1.7 variants, again supporting recombination rather than repeat mutation as the mechanism by which VOC-2020B were generated [46]

**S L452R** *-* This is a core change in the VOC-2021A (Table 1, Figure 4a-b). Although L452 does not interact directly with ACE2, L452R-carrying pseudovirus displayed increased infectivity *in vitro* [47], which could be due to a stronger binding to ACE2 via the electrostatic interaction with E35. However, L452R RBD expressed in yeast only slightly improved ACE2-binding [40]. Alternatively, the mutation may cause local conformational changes that impact the interactions of the RBD within the spike trimer or with ACE2. Notably, site 452 resides in a significant conformational epitope and L452R was shown to decrease binding to neutralizing antibodies (Figure 4b) [47, 48].

**S S477N** - The S S477N mutation spread rapidly in Australia (Figure 1, Figure 3b). Site 477, located at loop β4-5 of the RBD, is predicted not to establish persistent interactions with ACE2 [39]. Molecular dynamics simulations suggest that S477N affects the local flexibility of the RBD at the ACE2-binding interface, which could be underlying the highest binding affinity with ACE2 reported from potential mean force calculations and deep mutational scanning [40] [49]. Additionally, this site is located near an epitope and may alter antibody recognition and counteract the host immune response (Figure 4b).

**S H69_V70del** - The H69_V70del (in B.1.1.7) is adjacent to a linear epitope at the N-terminal domain of S (Figure 4a,c) [35], suggesting it may improve fitness by reducing host antibody effectiveness.

**S P681R and P681H** *-* Mutations in the multibasic furin cleavage site impacts cell-cell fusion [50, 51] and syncytia (i.e., multinucleate fused cells). Changes here may be a key factor underlying pathogenicity and virulence of SARS-CoV-2 strains [50, 52]. The P681R substitution present in B.1.617 exhibits a remarkable increase in syncytium formation in lung cells, which may explain the increased severity of the disease [23]. The similar substitution, P681H may have a comparable effect.

### 2.4. Potential adaptive mutations in non-S proteins

The potential epistatic partners of the S mutations are proposed to alter the viral replication resources and enhance antagonism to host innate immune responses (Table 2).

**N R203M** - The nucleocapsid (N) protein is the viral genome scaffold, is the most antigenic [53], and is necessary for viral replication [21, 54]. We previously reported that the Ser-Arg-rich motif of this protein (a.a. 183-206) displayed a high number of amino acid changes early in the pandemic consistent with it being under positive selection. A study using Oxford Nanopore^TM^ detected site-specific epigenetic modifications necessary for viral replication [26]. One of the epigenetic site resides in two highly successful SARS-CoV-2 haplotypes; a triple mutation at 28881-28883 (GGG to AAC, R203K, G204R) now found in nearly half of all sequences sampled (diamond nodes, Figure 1) and R203M, which is a defining mutation for B.1.617. This genomic region is highly conserved across several hundred years of coronavirus evolution (Figure S4) [27]. The R203K and G204R changes were recently shown to increase levels of the subgenomic RNA transcripts for N [55] and given that these epigenetic sites were discovered because the RNA pauses as it crosses the molecular nanopore sequencer, it may be that mutations here remove the epigenetic modification and speed the SARS-CoV-2 genome through the replisome. Interestingly, it has been proposed that the R203K, G204R double mutation arose through recombination [55].

**nsp2 I120F** - The role of the nonstructural protein 2 (nsp2) is unknown, but appears to involve multiple interactions with host proteins from a range of processes [56]. Deep learning-based structure prediction and cryo-electron microscopy density recently provided the atomic model of nsp2 (PDB ID 7MSW) and structural information was also used to localize the surfaces that are key for host protein-protein interactions with nsp2 [56]. From this it is hypothesized that nsp2 interacts with ribosomal RNA via a highly conserved zinc ribbon motif to unite ribosomes with the replication-transcription complexes.

The functional impact of the mutation I120F in nsp2 in the L-VOC D.2 is of interest given the MJN results. Site 120, identified in the nsp2 structure on Figure 6a, is a point of hydrophobic contact between a small helix, rich in positively charged residues, and a zinc binding site. The positively charged surface of the helix may be especially relevant for a putative interaction with the phosphate groups from ribosomal RNA. Normal mode analysis from DynaMut2 predicts that the substitution has a destabilizing effect in the protein structure (estimated ΔΔG^stability^ = -2 kcal/mol) [57]. Possibly, this could be caused by π-π stacking interactions of the tyrosine with aromatic residues in the same helix that would disrupt the contacts anchoring it to the protein core (Figure 6b). Additionally, site 120 is spatially close to E63 and E66, which were shown to be relevant for interactions with the endosomal/actin machinery via affinity purification mass spectrometry in HEK293T cells. Remarkably, upon mutation of these glutamates to lysines, there is increased interactions with proteins involved in ribosome biogenesis [56].

**Fig. 6.**
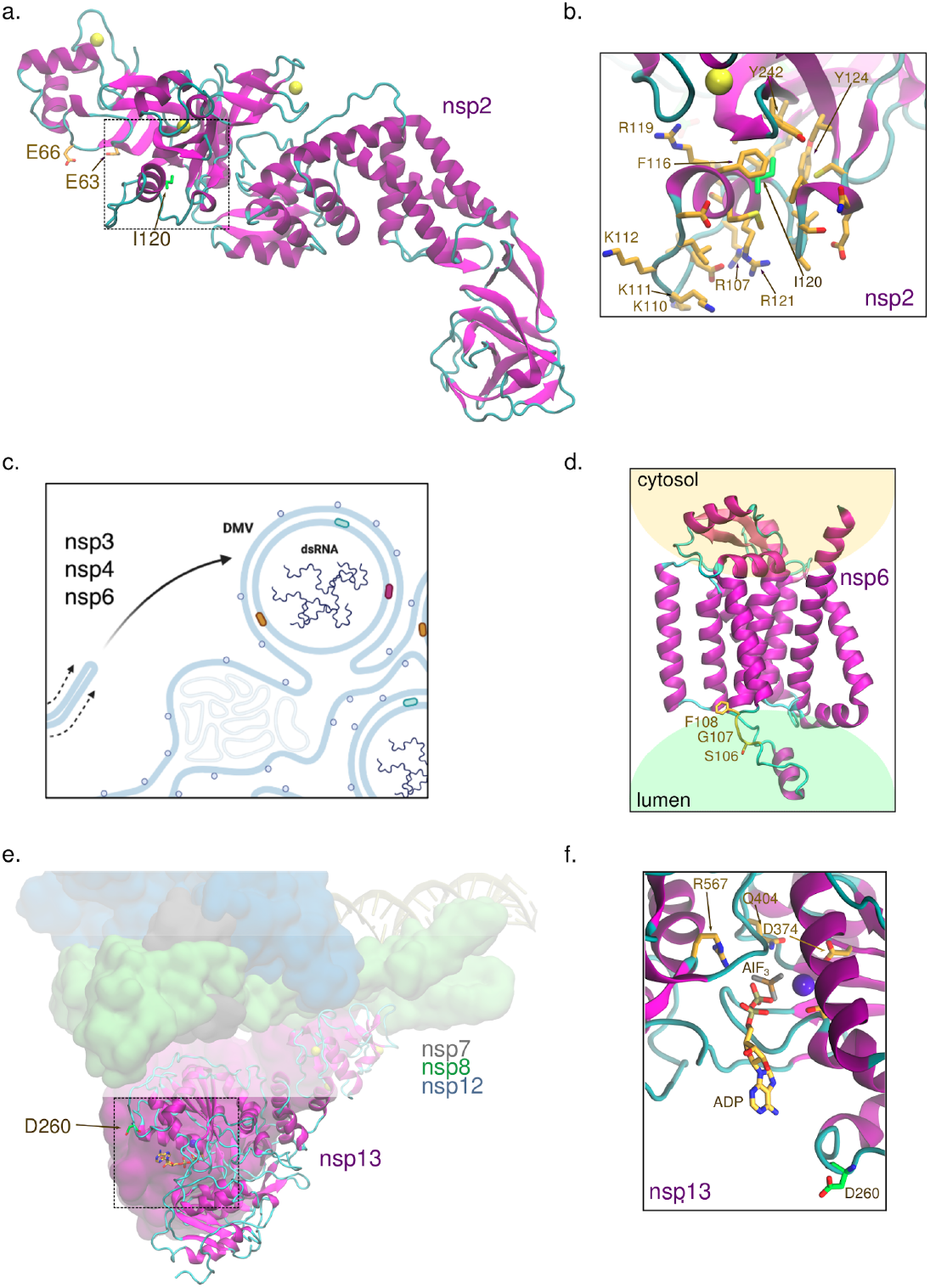
Location of mutations of prevalent SARS-CoV-2 variants on the structure of the nonstructural proteins nsp2, nsp6, and nsp13. a. Site 120 in nsp2 is located in a small helix near a zinc-binding site and residues E63 and E66, which play a role in the interaction with proteins involved in ribosome biogenesis and in the endosomal/actin machinery [56]. PDB ID 7MSW was used. b. I120 forms some of the hydrophobic contacts that anchor the helix at the surface of nsp2, where this site resides, to the protein core. c. Nsp6 participates in generating double-membrane vesicles (DMV) for viral genome replication. Natural selection for the biological traits of viral entry and replication may explain the increased transmission of variants with adaptive mutations in both S and nsp6. DMVs isolate the viral genome from host cell attack to provide for efficient genome and sub-genome replication and generate virions. d. Sites 106-108 are predicted to be located at/near the protein region of nsp6 embedded in the endoplasmic reticulum lumen (structure generated by AlphaFold2 [58]). e. Nsp13 is the SARS-CoV-2 helicase and it is part of the replication complex. f. D260 in nsp13 is mutated to tyrosine in B.1.427 and B.1.429 and it is located at the entrance of the NTP-binding site. PDB ID 6XEZ was used [59].

**nsp6 S106_F108del** - Nsp6 is critical for viral replication and suppression of the host immune response (Figure 5a and Figure 6c) [60]. Nsp3, nsp4, and nsp6 are responsible for producing double-membrane vesicles from the endoplasmic reticulum (ER) to protect the viral RNA from host attack and increase replication efficiency (Figure 6c) [61]. The nsp6 S106_F108del is predicted to be located at a loop in the interface between a transmembrane helix and the ER lumen based on a preliminary structural analysis by the AlphaFold2 system (Figure 6d), and we hypothesize that the deletion may affect functional interactions of nsp6 with other proteins. In addition, in agreement with the enhanced suppression of innate immune response reported for B.1.1.7 [62], changes in immune-antagonists, such as nsp6 S106_F108del, may be key to prolonged viral shedding [63].

**nsp13 D260Y** - Nsp13 or the helicase is a component of the replication-transcription complex that unwinds the duplex into single strands in a NTP-dependent manner [64]. The helicase and NTPase activities of nsp13 are highly coordinated, and mutations at the NTPase active site impair both ATP hydrolysis and the unwinding process [65]. The substitution D260Y, present in B.1.427 and B.1.429, is located at the entrance of the NTPase active site and may favor π-π stacking interactions with nucleobases (Figure 6e-f). Given that at high ATP concentrations, SARS-CoV nsp13 exhibits increased helicase activity on duplex RNA [66], it is possible that, similarly, the putative optimization on NPT uptake in nsp13 D260Y favors RNA unwinding.

Additionally, nsp13 is an important antagonist of the innate immune response (Figure 5a); it inhibits the type I interferon response by directly binding to TBK1 that impedes IRF3 phosphorylation [67]. The dual role of nsp6 and nsp13 in immune suppression and viral replication may suggest a convergent evolution of SARS-CoV-2 manifested in most of the VOC, which carries either nsp6 S106_F108del or nsp13 D260Y.

## 3. Concluding Remarks

Our network-based analysis of SARS-CoV-2 evolution indicates that mutations in S and in non-S proteins act in an epistatic manner to enhance viral fitness. Particularly, the S D614G substitution increases infectivity and is now predominant in the circulating virus [68], and S N501Y is associated with higher virulence [42]. We show that the expansion of the strains carrying these substitutions only occurred upon their combination with nsp12 L323P [21] and nsp6 S106_F108del, respectively. A hypothesis consistent with these observations is that the changes in S enhance viral entry into the host cells and better escape neutralizing antibodies, but they do not easily transmit due to rapid suppression by a robust innate immune response. A second mutation thus counteracts the immune-driven suppression. In the case of S D614G, the nsp12 L323P may have increased the replication rate of the virus to outcompete the immune response, which is supported by quantification of viral strains in clinical samples [21, 69]. However, the separate effects of S D614G and nsp12 L323P could not be ascribed in the referred study [69] because it did not include individuals infected with variants harboring only one of the mutations.

In the variants carrying S N501Y (VOC-2020B), we hypothesize that nsp6 S106_P108del may affect viral replication in DMVs or suppress the interferon-driven antiviral response [70]. Other mutations may also contribute to that. For example, it was recently shown *in vitro* that B.1.1.7 exhibits enhanced innate immune evasion, which was attributed to increased transcription of *orf9b*, nested within the nucleocapsid gene [62], although it was not ruled out that this was due to nsp6 S106_F108del.

Our structural analyses identify other mutations shared among different VOC that reside in key locations of proteins involved in viral replication and/or in innate immune antagonism, such as nsp13 D260Y, suggesting a convergent evolution of SARS-CoV-2 (Table 2). This emphasizes the importance of tracking mutations in a genome-wide manner as a strategy to avoid the emergence of future VOC. For example, an earlier dominant variant in Australia (D.2) that carried the mutations S477N in S and I120F in nsp2 was successfully restrained. However, variants harboring only the S S477N are currently circulating in several European countries (Table S5) and may originate the next VOC if combined with a functionally complementary mutation. We demonstrate that recombination events may be accelerating the junction of haplotypes carrying adaptive and cooperative mutations in S and in non-S proteins and that repeat mutations do not explain the high mutation load of VOC-2020B.

An equally significant outcome from the VOC-defining mutations is escape from the adaptive immune response [71] (Figure 5c). As a case in point, the resurgence of COVID-19 in Manaus, Brazil, in January 2021, where seroprevalence was above 75% in October 2020, is due to immune escape of new SARS-CoV-2 lineages [1]. Broad disease prevalence and community spread of COVID-19 increase the probability that divergent haplotypes come in contact resulting in adaptive epistatic mutations that dramatically enhance viral fitness. This emphasizes that regions with low sequence surveillance can be viral breeding grounds for the next SARS-CoV-2 VOC.

## Data Accessibility

All SARS-CoV-2 sequences used in this study are available from the public repositories Genome Initiative on Sharing All Influenza Data (GISAID, gisaid.org), the National Center for Biotechnology Information (NCBI, https://www.ncbi.nlm.nih.gov/sars-cov-2/) and the COVID-19 Genomics UK Consortium (COG, https://www.sanger.ac.uk/collaboration/covid-19-genomics-uk-cog-uk-consortium/

## Author Contribution

**MR Garvin**: Conceptualization, Data curation, Funding acquisition, Formal Analysis, Investigation, Methodology, Visualization, Writing – original draft, Writing – review & editing.

**ET Prates**: Formal Analysis, Investigation, Visualization, Writing - original draft, Writing - review & editing

**J Romero**: Conceptualization, Formal Analysis, Investigation, Methodology, Software, Writing – original draft, Writing – review & editing.

**A Cliff**: Methodology, Software, Writing – review & editing

**JGFM Gazolla**: Software, Formal Analysis, Investigation, Data Curation, Visualization, Writing - Review and Editing.

**M Pickholz:** Investigation, Visualization, Writing – original draft, Writing – review & editing

**M Pavicic:** Investigation, Writing – original draft, Writing – review & editing

**DA Jacobson**: Conceptualization, Funding acquisition, Formal Analysis, Investigation, Project administration, Supervision, Resources,Writing – original draft, Writing – review & editing

## Acknowledgments

The viral evolution research was funded by the Laboratory Directed Research and Development Program of Oak Ridge National Laboratory, managed by UT-Battelle, LLC for the US Department of Energy (LOIS:10074) and the structural implication work was funded via the DOE Office of Science through the National Virtual Biotechnology Laboratory (NVBL), a consortium of DOE national laboratories focused on the response to COVID-19, with funding provided by the Coronavirus CARES Act. This work was also funded by the United States Government. This research used resources of the Oak Ridge Leadership Computing Facility (OLCF) and the Compute and Data Environment for Science (CADES) at the Oak Ridge National Laboratory, which is supported by the Office of Science of the U.S. Department of Energy under Contract No. DE-AC05-00OR22725. Figures generated with Biorender and VMD. We gratefully acknowledge the Originating laboratories responsible for obtaining the viral specimens and the Submitting laboratories where genetic sequence data were generated and shared via the GISAID Initiative, on which this research is based.

## Box 1.

### Networks versus phylogenetic trees for studies of SARS-CoV-2

The COVID-19 pandemic is an unprecedented tragedy but also an opportunity to study molecular evolution given the global sampling of the mutational space of SARS-CoV-2. The majority of current efforts analyzing evolution employ phylogenetic trees, which are useful for studying species, but cannot effectively incorporate critical information needed to study populations of an emerging pathogen such as this [11, 12]. Indeed, a recent study found that coronavirus evolution shows very little resemblance to a tree structure [32], and therefore it is pertinent to ask not if a phylogenetic tree is useful for studying the evolution of SARS-CoV-2, but rather, what is the best tool to do so. In evolutionary analyses of species, a single sequence represents each taxon and variable sites are assumed to be fixed within a species; variation across species therefore represents an estimate of evolutionary substitution rate across time, typically measured in thousands or millions of years and focused on amino acid changes. Trees are typically rooted (an origin is provided) and directed (evolution proceeds from root to tip) and therefore estimates of evolutionary events (e.g. speciation) can be derived. Tips of trees are usually the unit of interest and internal nodes are largely ignored.

In contrast, for populations, mutations are *segregating polymorphisms* that occur at different frequencies in different locations at different times and, in the case of SARS-CoV-2, evolution is often measured in days, weeks, or months. A haplotype network is ideal to study populations such as SARS-CoV-2 for several reasons. Firstly, it assumes single, ordered mutational steps where each node represents a haploid RNA sequence (haplotype) and the edge between nodes is a mutation or mutations leading to a new one. Using a median-joining-network (MJN) approach, extant haplotypes for a taxon are sampled whereas unsampled, or extinct lineages are inferred. Here, SARS-CoV-2 sequence data repositories provide extensive sampling of haplotypes and collection dates (the calendar date of the sample). Given that the temporal distribution of haplotypes is inherent in an MJN (the model assumes time-ordered mutations), the mutational history of the virus can be traced as a genealogy that can incorporate both the relative *and* absolute time. Haplotype networks are not implicitly rooted or directed, but in the case of SARS-CoV-2, we have a root (Wuhan reference from 2019) and sampling times provide direction. Internal nodes and tips are both inherent in the model.

Secondly, important metadata that are relevant to populations but not species such as frequency, date of emergence, mutations of interest, effective reproduction number, or clinical outcomes [72] can simultaneously be displayed on the haplotype network but not the phylogenetic tree, especially for hundreds of thousands to millions of sequences as is the case with SARS-CoV-2. Third, and perhaps most importantly, when the MJN model fails, it produces loops or clusters of inferred haplotypes on a network that can indicate recombination events, back mutations, or repeat mutations at a site. In contrast, a phylogenetic tree algorithm forces a bifurcating structure that cannot indicate if these events have occurred.

To demonstrate, here we generated a network and a phylogenetic tree of SARS-CoV-2 haplotypes sampled during the first four months of the pandemic from GISAID (A Global Initiative on Sharing All Influenza Data). Early in the evolution of SARS-CoV-2, phylogenetic trees and haplotype network indicate similar topologies, but clearly, frequency, emergence date, temporal order of adaptive mutations, and other important metadata can easily be displayed simultaneously on a network, but not a tree. At day 96, reticulations (homoplasy loops) begin to appear in the network, indicating reverse mutations to the ancestral states or possibly recombination events. Polytomies (molecular events often found in rapidly expanding populations) are another important feature identified when using networks, but are lost when using phylogenetic trees. For example, haplotype H04 in the MJN-derived network represents a hard polytomy and indicates that a frequent variant is further undergoing multiple independent mutational events, but the phylogenetic tree is unable to convey this information, mainly due to its focus on tips and not internal nodes.

**Fig. 7.**
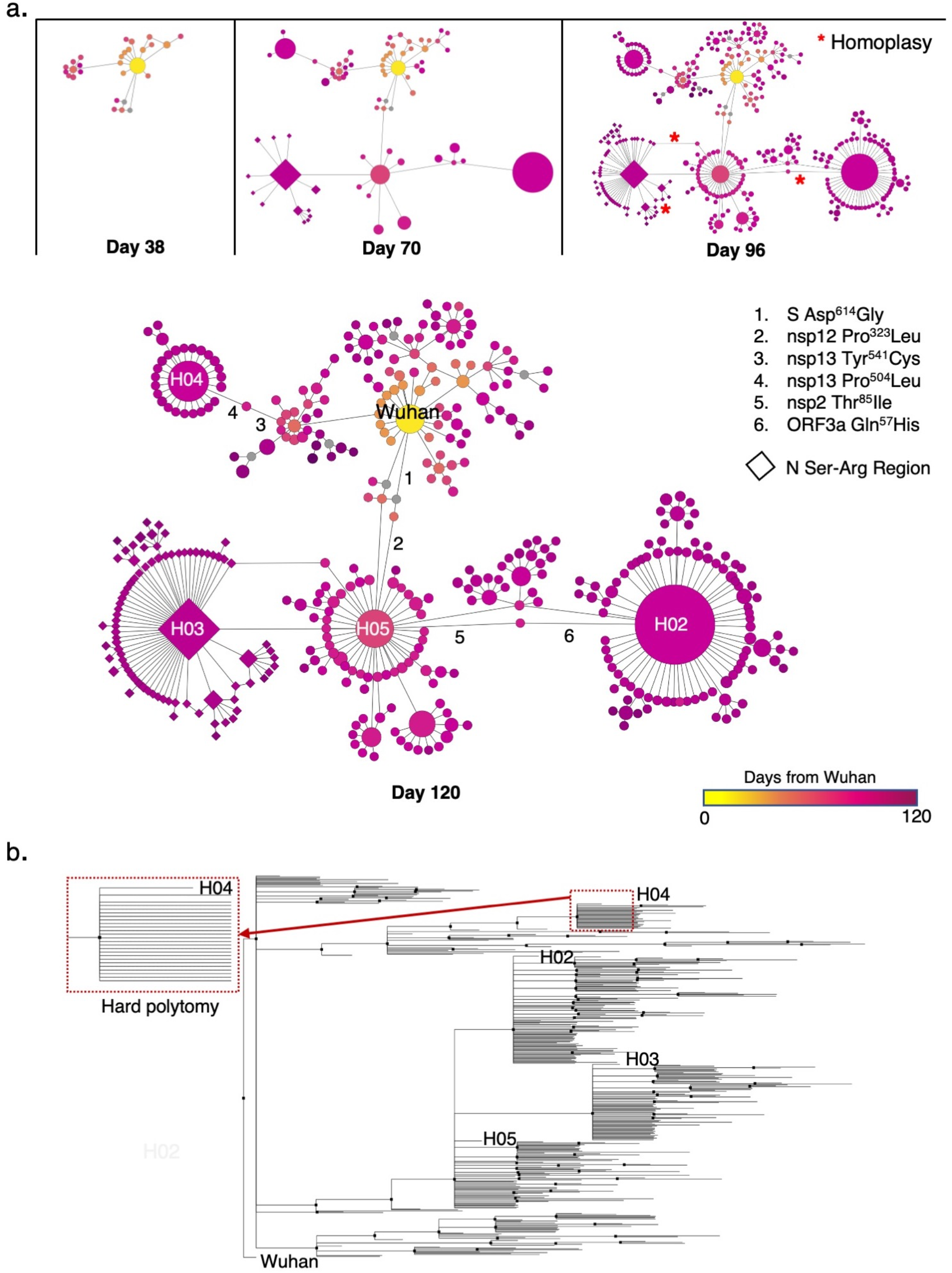
Comparison of a haplotype network and phylogenetic tree generated with SARS-CoV-2 sequences sampled through April 2020. **a**. MJN-derived haplotype network of SARS-CoV-2 at 38, 70, 96, and 120 days. Node sizes in the MJN correspond to sample sizes for a given haplotype and node colors indicate the time of its first report relative to the putative origin of the pandemic in Wuhan. Gray nodes are inferred haplotypes. The most abundant haplotypes are named H02 - H05 and numerals 1 - 6 identify several mutations discussed here and in our previous work [21]. Diamond shape nodes denote haplotypes that harbor a 3 nucleotide mutation in the nucleocapsid gene (N) that is highly conserved and directly affects viral replication *in vitro* [54, 62]. **b.** The phylogenetic tree is unable to convey the same information. For example, rapidly expanding populations often display polytomies, i.e., single mutations from a common central haplotype. Those events are readily identified on the haplotype network, but difficult to interpret on a tree because they are usually visualized as a multi-pronged fork (outlined in the dashed-line box) rather than a star pattern (compare H04 in (a) and (b)). These true biological processes also cause tree algorithms to perform poorly because they violate their assumptions, slowing convergence. Additionally, MJN-derived haplotype networks are able to indicate reticulations (i.e., loops) that could denote recombination, reverse mutations, or other biologically important events whereas the forced bifurcation of phylogenetic tree algorithms is unable to display these. Reference sequence: NC_045512, Wuhan, December 24, 2019.

## Methods

### Sequence data pre-processing

We downloaded SARS-CoV-2 sequences in FASTA format and corresponding metadata from GISAID and processed as we have reported previously [21, 73]. To ensure that deletions were accounted for, full genome sequences were aligned with MAFFT [74] to the established reference genome (accession NC_045512), uploaded into CLC Genomics Workbench, and trimmed to the start and stop codons (nsp1 start site and ORF10 stop codon). Aligned sequences in tab-delimited format were imported into R to count the number of variable accessions at each of the 29,409 sites.

Variable sites were determined with all sequences downloaded up through the end of January, 2021. In order to reduce false-positive mutation sites (those that were due to technical error), we selected sites that were variable in 25 or more individuals (0.01%) compared to the reference (all 25 were required to be the same state: A, G, T, C, or -). We further pruned these by removing sites in which 20% or more of the accessions harbored an unknown character state (“N”), leaving 2,128 variable sites for downstream analyses. After removing sequences with an “N” at any of these sites, we retained 280,409 individuals. Prior to submission, we updated the number of sequences through April 19, 2021, keeping the same 2128 variable sites, which allowed us to capture the most up-to-date metadata and produced 640,211 for analysis. We kept haplotypes that occurred in more than 35 individuals to remove rare or artifact-derived haplotypes (https://virological.org/t/issues-with-sars-cov-2-sequencing-data/473).

For the comparison of median-joining networks and phylogenetic trees, we used sequences from the pandemic sampled through the end of April, 2020. We used variable sites found in more than ten individuals and haplotypes found in five or more individuals as we had in previous work [21]. This produced 410 unique haplotypes based on 467 variable sites.

### Median-joining network (MJN)

Haplotypes were coded in NEXUS format and uploaded to PopArt [75]. An MJN was produced with the epsilon parameter set to 0. The networks were exported as a table and visualized in Cytoscape [76] with corresponding metadata. The date of emergence of each haplotype was defined by the sample date subtracted from the report date for the Wuhan reference sequence (December 24, 2019) and then one day was added to remove zeros. For samples that only reported the month but no day, we recorded the day as the 15th of that month. We excluded samples with no sampling date.

### Phylogenetic tree

We used the program MrBayes to generate a phylogenetic tree [77]. Parameters were set to *Nucmodel=4by4*, *Nst=6*, *Code=Universal*, and *Rates=Invgamma*. We performed 5,000,000 mcmc generations, which produced a stable standard deviation of split frequencies of 0.014. A consensus tree was generated using the 50% majority rule and visualized using FigTree v1.4.4 (http://tree.bio.ed.ac.uk/software/figtree/).

### Estimation of genome mutation load

We estimated the mutation load using two data sets. First, we used the 640,211 sequences based on 2,128 variable sites used for the MJN because these represent high-confidence mutations. For each of the 640,211 accessions, we counted the number of differences of the 2,128 variable sites compared to the reference genome (accession NC_045512) and recorded the day of emergence. The mutational load for all accessions for a given day was then averaged and this was plotted across time. For the second estimate of mutation rate, we used all variable sites across the full genome (29,409 sites) to include rare variants and removed all sequences with at least one ambiguous site, leaving 584,119 accessions.

For the population-level estimate of mutation accumulation, we applied the filters used to identify the 2,128 variable sites that were used for the MJN for all sequences up through April 19, 2021. We did not include new mutations because the B.1.1.7 VOC and its downstream haplotypes had become the predominant variants globally at that time and, consequently, much early information of the molecular evolution is lost when applying frequency filters on the entire GISAID database. This is exacerbated with the MJN approach because the software algorithm used to generate the network is computationally intractable with greater than 1,000 haplotypes and therefore future efforts will either need to ignore early molecular events or use new methods that can handle the large datasets and any recombination events that occur (an alternative approach would be to now use the alpha or delta variant as the reference sequence because they are now the predominant strains globally).

For calculations of population-level mutation accumulation, it is possible (and necessary) to include all sequences to determine if mutation or recombination are the cause of the high mutation load seen in the late 2020 VOC. After applying the frequency and haplotype filters, we retained 5,011 variable sites that define 12,282 unique haplotypes for further analysis. Mutations to five possible states (A, G, T, C, and -) were counted at each site on the first date that they appeared and their appearance at later dates were excluded. Multiple mutations at a site to different states were counted with this method.

For lineage-specific mutation curves, we extracted all sequences based on their PANGO lineage listed in the metadata from GISAID that also had a sample data and plotted the cumulative number over time, where time is represented by days from first appearance. To estimate the rate of accumulation, we calculated the slope for the linear portion of each of the curves.

### Probability of mutation accumulation

To calculate the chance of accumulating several mutations in a certain period, the probability density function for a normal distribution is used:

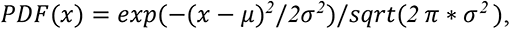

where *μ* is the expected number of mutations for that date, *x* is the measured value, and *σ* is the standard deviation of error calculated from the data shown in Fig. 1b, considering the difference between the actual and predicted number of mutations . The expected value of mutations *μ* for a given time period is computed from the estimated rate of mutations per day (Figure 3, 0.05). c. The period of interest to our discussion (June-October 2020) corresponds to 122 days, for which, the integral of PDF(*x*=13) gives the probability of 1*10^-15^ to accumulate 13 mutational events.

### Screen for coinfected individuals with UK B.1.1.7

We extracted 25 samples from the Sequence Read Archive at NCBI for each of the months of October, November, December, and January listed as variant B.1.1.7 from the UK (Table S2) for a total of 100 samples to check for coinfection. The reads were mapped to the NC_045512 Wuhan reference using CLC Genomics Workbench using the default parameters except for length fraction and similarity fraction were set to 0.9. Three sites specific to UK B.1.1.7 were analyzed for possible heterozygosity. Of the 100 we sampled, two appeared to be cases of coinfection. This supports the hypothesis that the large expansion in overall mutations seen in UK B.1.1.7 are likely due to recombination. In addition, it also supports the case that coinfection is occurring at a baseline sufficient to allow for occasional recombination.

### Protein structure analysis

VMD was used to visualize the protein structures and analyze the potential functional effects of mutations [78]. Figure 3 was created using Inkscape (https://inskape.org/) and Gimp 2.8

(https://www.gimp.org) [79].

### Molecular dynamics simulations

Molecular dynamics (MD) simulations were used to study interactions between SARS-CoV-2 RBD and ACE2 from ferret and human. Three independent extensive MD simulations were performed for each species using GROMACS 2020 package [80] and the CHARMM36 force field for protein and glycans [81, 82]. Each simulation ran up to 800 ns, being the last 500 ns used for analysis. PDB id 6M17 was used to build the ACE2-RBD complexes. Given the high sequence identity between human and ferret ACE2 (83%), we performed local modeling of the non-conserved amino acid residues in ferret ACE2 using the human homolog as the template, via RosettaRemodel [83].

The inputs for simulations were generated using CHARMM-GUI [84]. Counterions were added for electroneutrality (0.1 M NaCl). The complexes were surrounded by TIP3P water molecules to form a layer of at least 10 Å relative to the box borders [85]. Simulations were performed using the NPT ensemble. The temperature was maintained at 310 K with the Nosé–Hoover thermostat using a time constant of 1.0 ps [86]. The pressure was maintained at 1 bar with the isotropic Parrinello–Rahman barostat using a compressibility of 4.5 × 10^−5^ bar^−1^ and a time constant of 1.0 ps in a rectangular simulation box [87]. The particle mesh Ewald method was used for the treatment of periodic electrostatic interactions with a cutoff distance of 1.2 nm [88]. The Lennard–Jones potential was smoothed over the cutoff range of 1.0–1.2 nm by using the force-based switching function. Only atoms in the Verlet pair list with a cutoff range reassigned every 20 steps were considered. The LINCS algorithm was used to constrain all bonds involving hydrogen atoms to allow the use of a 2 fs time step [89]. The suggested protocol for nonbonded interactions with the CHARMM36 force field when used in the GROMACS suite was followed.

The Hbonds plugin in VMD was used to identify hydrogen bond interactions along the simulations [78]. The geometric criteria adopted are a cutoff of 3.5 Å for donor-acceptor distance and 30° for acceptor-donor-H angle. The Timeline plugin was used to count contacts formed by a given amino acid residue. We defined the distance of 4 Å between any atom pairs as the cutoff for contact.

---

**Fig. S1.**
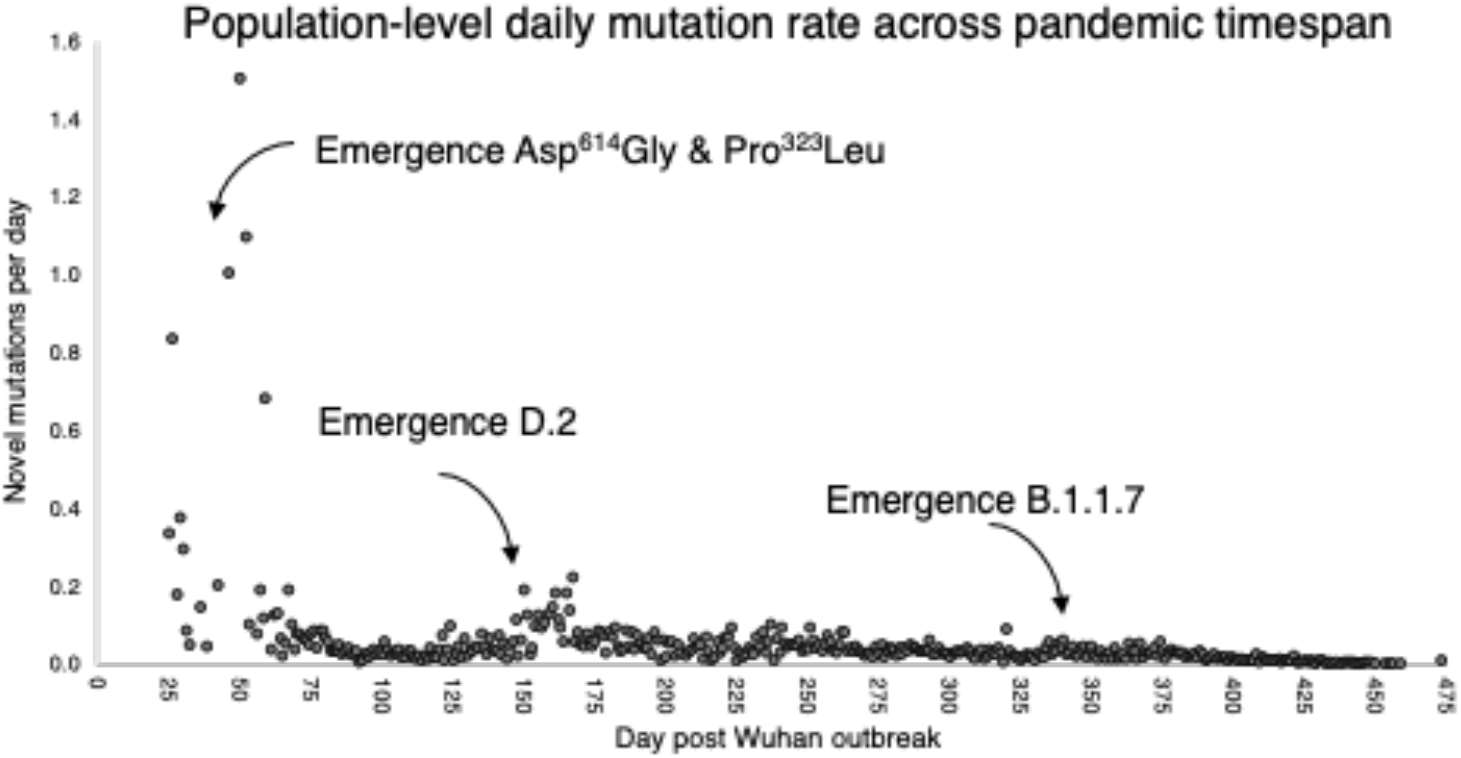
Population level mutation rate over the course of the pandemic. Number of novel mutations sampled across the globe for each day are plotted against time (days from the Wuhan outbreak). Emergence of major VOC are provided for context and show small increases in the number of new mutations but there is an overall decrease across time, even accounting for multiple mutations at a site to different nucleotide states and deletions.

**Fig. S2.**
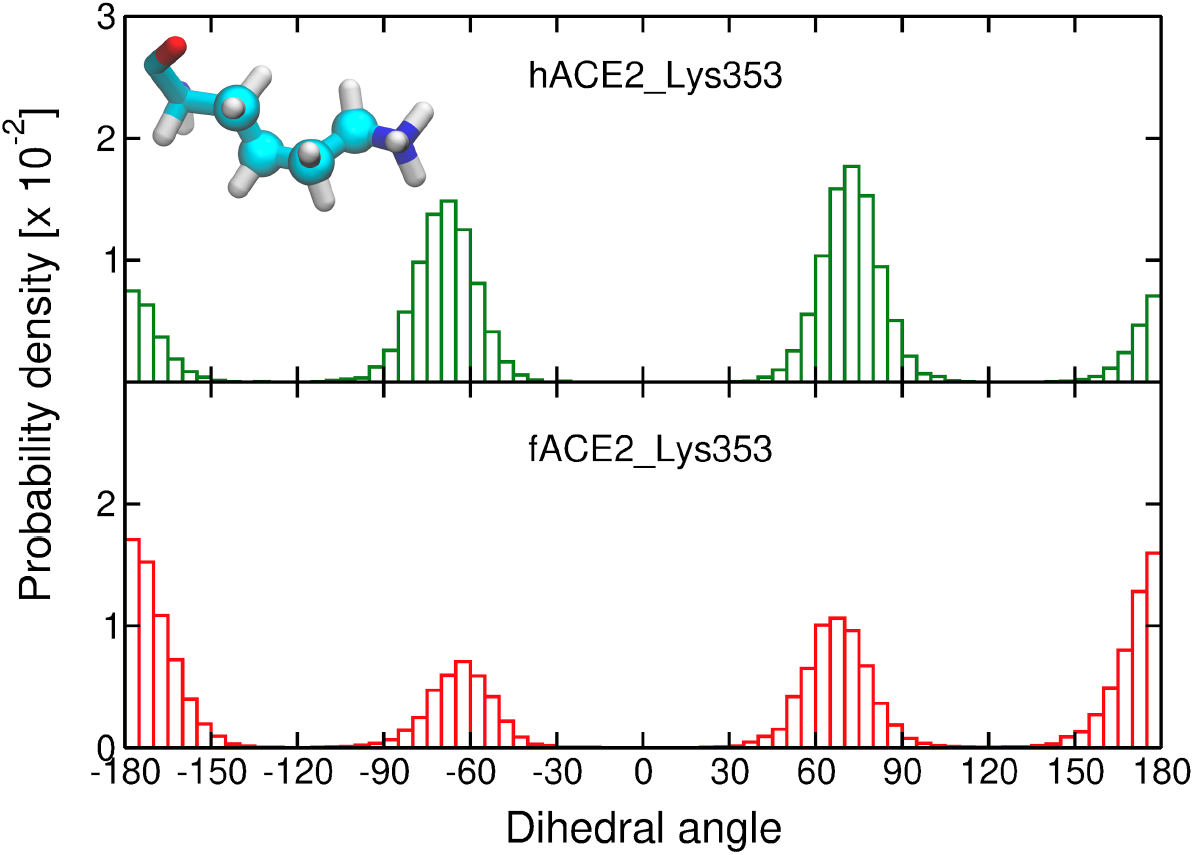
The probability density of the conformations of Lys^353^ in human and ferret ACE2 in the simulations. Histograms of the distribution of a dihedral angle of the Lys^353^ side chain carbon atoms in human ACE2 (hACE2, upper figure) and ferret ACE2 (fACE2, lower figure) in complex with the SARS-CoV-2 S receptor-binding domain. The atoms forming the selected dihedral are depicted as spheres in the molecular representation of Lys^353^. Three independent simulations are considered for the calculation of the histograms. Dihedral angles near ±180° correspond to a more stretched conformation (i.e., *trans*).

**Fig. S3.**
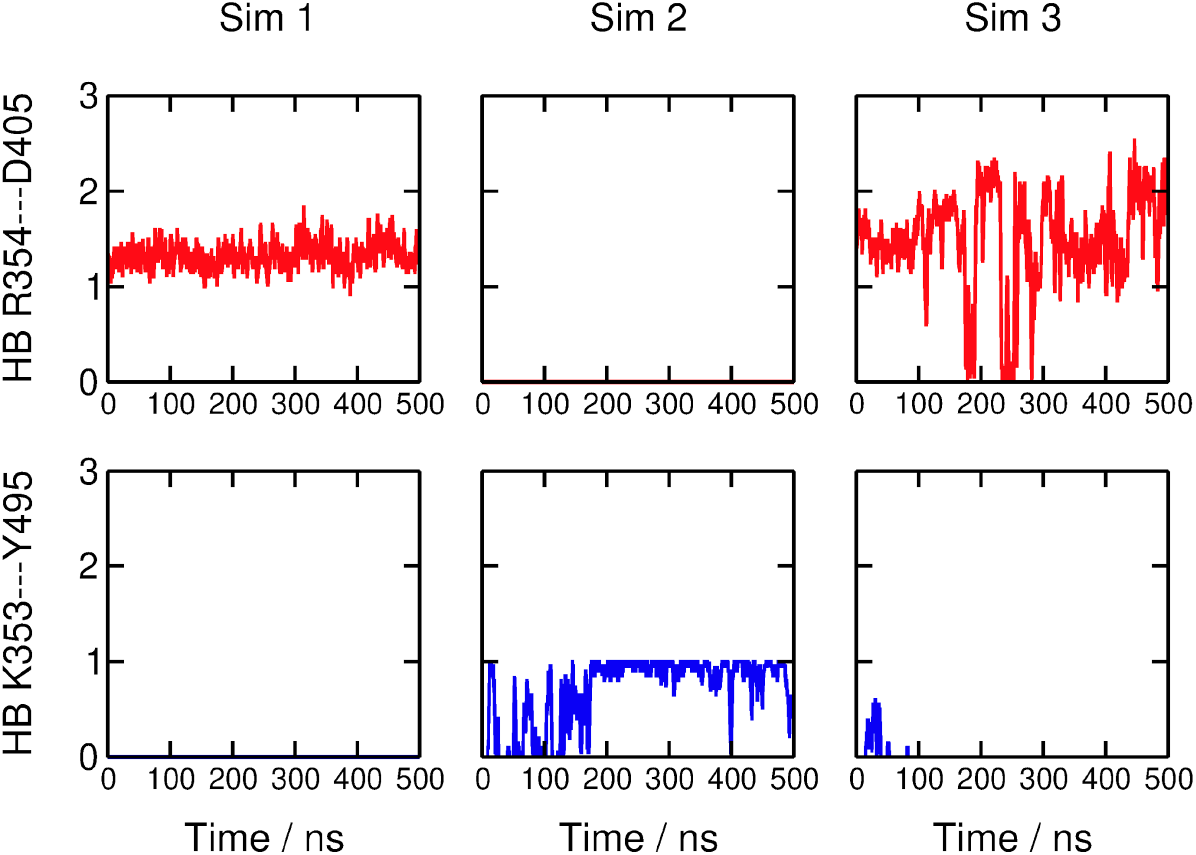
Competing hydrogen bond interactions formed between positively charged amino acid residues in ferret ACE2 (fACE2) and the SARS-CoV-2 S receptor-binding domain. Time evolution of the number of hydrogen bonds (HB) that fACE2 Arg3^54^ and Lys^353^ form with Asp^405^ and Tyr^495^ from the SARS-CoV-2 S receptor-binding domain. The columns correspond to the three simulation replicas. The geometric criteria adopted for hydrogen bonds are a cutoff of 3.0 Å for donor-acceptor distance and 20° for acceptor-donor-H angle.

**Fig. S4.**
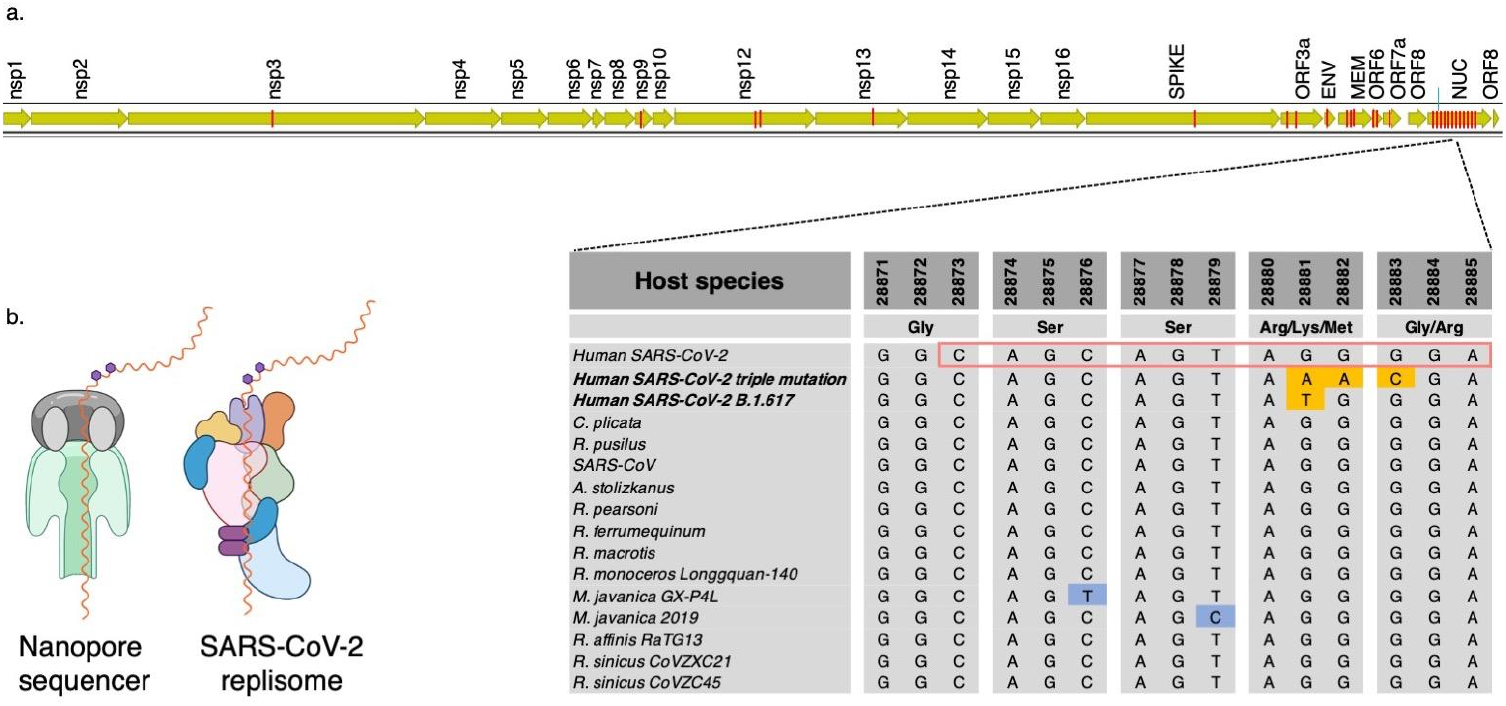
Modifications at the Ser-Arg-rich region of N may affect replication speed. **a.** Location of 41 epigenetic sites reported in Kim et al. 2020 (red bars on SARS-CoV-2 genome). One of the sites in the nucleocapsid gene (nucleotides in red box of aligned sequences) is highly conserved across diverse host-defined coronaviruses. All bats and human coronavirus species from China are completely conserved at the epigenetic site 28881-28883, except for a 3-bp mutation in SARS-CoV-2 that occurred early in the pandemic and now corresponds to ∼50% of all sequences globally (diamond nodes in Figure 1). **b.** Kim et al. proposed that *N^6^*-methyladeonsine modification of the genome (purple hexagons), common in RNA viruses, caused the strand to pause while traversing the nanopore sequencing apparatus. We propose that loss of this site via mutations at site 203 in N may increase the replication rate of the RNA strand through the SARS-CoV-2 replisome. *Aselliscus stoliczkanus* - Stoliczka’s trident bat, *Chaerephon plicata* - wrinkle-lipped free-tailed bat, *Rhinolophus pusillus -* least horseshoe bat, *R. pearsoni* - Pearson’s horseshoe bat*, R. macrotis* - big-eared horseshoe bat*, R. ferrumequinum* - greater horseshoe bat*, R. monoceros* - Formosan lesser horseshoe bat*, R. affinis* intermediate horseshoe bat*, R. sinicus* Chinese rufous horseshoe bat, *R. mayalanis* - Mayalan horseshoe bat, *SARS* - Severe Acute Respiratory Syndrome, *Manis javanica* - Malayan pangolin. *Created with BioRender.com*.

**Table S4.** Average number of contacts formed between Asn^501^ in the receptor-binding domains of SARS-CoV-2 S and residues in ACE2 from human (hACE2) and ferret (fACE2). A distance of 4 Å between any atom pairs was defined as the cut-off for contact statistics.

| ACE2 residue | hACE2\| | fACE2 |
| --- | --- | --- |
| Tyr<sup>41</sup> | 0.96 ± 0.02 | 0.80 ± 0.03 |
| Lys<sup>353</sup> | 0.99 ± 0.01 | 0.90 ± 0.01 |
| Asp<sup>353</sup> | 0.98 ± 0.01 | 0.70 ± 0.04 |

## Notes

### Competing Interest Statement

The authors have declared no competing interest.

### Summary of Updates

We incorporated information from a recent analysis that identified recombination in North America and revised Figure 3.

## References

1. Sabino EC, Buss LF, Carvalho MPS, Prete CA Jr, Crispim MAE, Fraiji NA, et al. Resurgence of COVID-19 in Manaus, Brazil, despite high seroprevalence. Lancet 2021.

2. Volz E, Mishra S, Chand M, Barrett JC, Johnson R, Geidelberg L, et al. Transmission of SARS-CoV-2 Lineage B.1.1.7 in England: Insights from linking epidemiological and genetic data.

3. Davies NG, Jarvis CI, John Edmunds W, Jewell NP, Diaz-Ordaz K, Keogh RH, et al. Increased mortality in community-tested cases of SARS-CoV-2 lineage B.1.1.7.

4. Alpert T, Brito AF, Lasek-Nesselquist E, Rothman J, Valesano AL, MacKay MJ, et al. Early introductions and transmission of SARS-CoV-2 variant B.1.1.7 in the United States. Cell 2021; 184: 2595–2604.e13.

5. Washington NL, Gangavarapu K, Zeller M, Bolze A, Cirulli ET, Schiabor Barrett KM, et al. Emergence and rapid transmission of SARS-CoV-2 B.1.1.7 in the United States. Cell 2021; 184: 2587–2594.e7.

6. Challen R, Brooks-Pollock E, Read JM, Dyson L, Tsaneva-Atanasova K, Danon L. Risk of mortality in patients infected with SARS-CoV-2 variant of concern 202012/1: matched cohort study. BMJ 2021; 372: n579.

7. Faria NR, Mellan TA, Whittaker C, Claro IM, Candido D da S, Mishra S, et al. Genomics and epidemiology of the P.1 SARS-CoV-2 lineage in Manaus, Brazil. Science 2021; 372: 815–821.

8. Funk T, Pharris A, Spiteri G, Bundle N, Melidou A, Carr M, et al. Characteristics of SARS-CoV-2 variants of concern B.1.1.7, B.1.351 or P.1: data from seven EU/EEA countries, weeks 38/2020 to 10/2021. Euro Surveill 2021; 26.

9. Tao K, Tzou PL, Nouhin J, Gupta RK, de Oliveira T, Kosakovsky Pond SL, et al. The biological and clinical significance of emerging SARS-CoV-2 variants. Nature Reviews Genetics . 2021., 22: 757–773

10. Singh J, Rahman SA, Ehtesham NZ, Hira S, Hasnain SE. SARS-CoV-2 variants of concern are emerging in India. Nat Med 2021.

11. Velasco JD. Phylogeny as population history. Philosophy and Theory in Biology . 2013., 5

12. Huson DH, Bryant D. Application of phylogenetic networks in evolutionary studies. Mol Biol Evol 2006; 23: 254–267.

13. Paradis E. Analysis of haplotype networks: The randomized minimum spanning tree method. Methods in Ecology and Evolution . 2018., 9: 1308–1317

14. Martin DP, Weaver S, Tegally H, San EJ, Shank SD, Wilkinson E, et al. The emergence and ongoing convergent evolution of the N501Y lineages coincides with a major global shift in the SARS-CoV-2 selective landscape. medRxiv 2021.

15. Bentley K, Evans DJ. Mechanisms and consequences of positive-strand RNA virus recombination. J Gen Virol 2018; 99: 1345–1356.

16. Simon-Loriere E, Holmes EC. Why do RNA viruses recombine? Nat Rev Microbiol 2011; 9: 617–626.

17. Zella D, Giovanetti M, Benedetti F, Unali F, Spoto S, Guarino M, et al. The variants question: What is the problem? J Med Virol 2021; 93: 6479–6485.

18. Gutierrez B, Castelán Sánchez HG, da Silva Candido D, Jackson B, Fleishon S, Ruis C, et al. Emergence and widespread circulation of a recombinant SARS-CoV-2 lineage in North America. medRxiv 2021; 2021.11.19.21266601.

19. Jackson B, Boni MF, Bull MJ, Colleran A, Colquhoun RM, Darby AC, et al. Generation and transmission of interlineage recombinants in the SARS-CoV-2 pandemic. Cell 2021; 184: 5179–5188.e8.

20. Benedetti F, Pachetti M, Marini B, Ippodrino R, Ciccozzi M, Zella D. SARS-CoV-2: March toward adaptation. J Med Virol 2020; 92: 2274–2276.

21. Garvin MR, Prates ET, Pavicic M, Jones P, Amos BK, Geiger A, et al. Potentially adaptive SARS-CoV-2 mutations discovered with novel spatiotemporal and explainable-AI models. Genome Biol 2020; in press.

22. Prates ET, Garvin MR, Pavicic M, Jones P, Shah M, Demerdash O, et al. Potential pathogenicity determinants identified from structural proteomics of SARS-CoV and SARS-CoV-2. Mol Biol Evol 2020.

23. Sheikh A, McMenamin J, Taylor B, Robertson C. SARS-CoV-2 Delta VOC in Scotland: demographics, risk of hospital admission, and vaccine effectiveness. The Lancet . 2021.

24. Muller HJ. THE RELATION OF RECOMBINATION TO MUTATIONAL ADVANCE. Mutat Res 1964; 106: 2–9.

25. Moutouh L, Corbeil J, Richman DD. Recombination leads to the rapid emergence of HIV-1 dually resistant mutants under selective drug pressure. Proc Natl Acad Sci U S A 1996; 93: 6106–6111.

26. Kim D, Lee J-Y, Yang J-S, Kim JW, Kim VN, Chang H. The Architecture of SARS-CoV-2 Transcriptome. Cell 2020; 181: 914–921.e10.

27. Boni MF, Lemey P, Jiang X, Lam TT-Y, Perry BW, Castoe TA, et al. Evolutionary origins of the SARS-CoV-2 sarbecovirus lineage responsible for the COVID-19 pandemic. Nat Microbiol 2020; 5: 1408–1417.

28. Gribble J, Stevens LJ, Agostini ML, Anderson-Daniels J, Chappell JD, Lu X, et al. The coronavirus proofreading exoribonuclease mediates extensive viral recombination. PLoS Pathog 2021; 17: e1009226.

29. Sender R, Bar-On YM, Gleizer S, Bernshtein B, Flamholz A, Phillips R, et al. The total number and mass of SARS-CoV-2 virions. Proc Natl Acad Sci U S A 2021; 118.

30. Peacock TP, Penrice-Randal R, Hiscox JA, Barclay WS. SARS-CoV-2 one year on: evidence for ongoing viral adaptation. J Gen Virol 2021; 102.

31. Anisimova M, Nielsen R, Yang Z. Effect of recombination on the accuracy of the likelihood method for detecting positive selection at amino acid sites. Genetics 2003; 164: 1229–1236.

32. Müller NF, Kistler KE, Bedford T. Recombination patterns in coronaviruses. bioRxiv 2021.

33. Wrapp D, Wang N, Corbett KS, Goldsmith JA, Hsieh C-L, Abiona O, et al. Cryo-EM structure of the 2019-nCoV spike in the prefusion conformation. Science 2020; 367: 1260–1263.

34. Casalino L, Gaieb Z, Goldsmith JA, Hjorth CK, Dommer AC, Harbison AM, et al. Beyond Shielding: The Roles of Glycans in the SARS-CoV-2 Spike Protein. ACS Cent Sci 2020; 6: 1722–1734.

35. Zhang B-Z, Hu Y-F, Chen L-L, Yau T, Tong Y-G, Hu J-C, et al. Mining of epitopes on spike protein of SARS-CoV-2 from COVID-19 patients. Cell Res 2020; 30: 702–704.

36. Yan R, Zhang Y, Li Y, Xia L, Guo Y, Zhou Q. Structural basis for the recognition of SARS-CoV-2 by full-length human ACE2. Science 2020; 367: 1444–1448.

37. Lauring AS, Hodcroft EB. Genetic Variants of SARS-CoV-2-What Do They Mean? JAMA 2021; 325: 529–531.

38. Shang J, Ye G, Shi K, Wan Y, Luo C, Aihara H, et al. Structural basis of receptor recognition by SARS-CoV-2. Nature 2020; 581: 221–224.

39. Ali A, Vijayan R. Dynamics of the ACE2-SARS-CoV-2/SARS-CoV spike protein interface reveal unique mechanisms. Sci Rep 2020; 10: 14214.

40. Starr TN, Greaney AJ, Hilton SK, Crawford KHD, Navarro MJ, Bowen JE, et al. Deep mutational scanning of SARS-CoV-2 receptor binding domain reveals constraints on folding and ACE2 binding.

41. Wang Z, Schmidt F, Weisblum Y, Muecksch F, Barnes CO, Finkin S, et al. mRNA vaccine-elicited antibodies to SARS-CoV-2 and circulating variants. Nature 2021.

42. Gu H, Chen Q, Yang G, He L, Fan H, Deng Y-Q, et al. Adaptation of SARS-CoV-2 in BALB/c mice for testing vaccine efficacy. Science 2020; 369: 1603–1607.

43. Oude Munnink BB, Sikkema RS, Nieuwenhuijse DF, Molenaar RJ, Munger E, Molenkamp R, et al. Transmission of SARS-CoV-2 on mink farms between humans and mink and back to humans. Science 2021; 371: 172–177.

44. Sawatzki K, Hill NJ, Puryear WB, Foss AD, Stone JJ, Runstadler JA. Host barriers to SARS-CoV-2 demonstrated by ferrets in a high-exposure domestic setting. Proc Natl Acad Sci U S A 2021; 118.

45. Richard M, Kok A, de Meulder D, Bestebroer TM, Lamers MM, Okba NMA, et al. SARS-CoV-2 is transmitted via contact and via the air between ferrets. Nat Commun 2020; 11: 3496.

46. Grabowski F, Preibisch G, Giziński S, Kochańczyk M, Lipniacki T. SARS-CoV-2 Variant of Concern 202012/01 Has about Twofold Replicative Advantage and Acquires Concerning Mutations. Viruses 2021; 13.

47. Deng X, Garcia-Knight MA, Khalid MM, Servellita V, Wang C, Morris MK, et al. Transmission, infectivity, and neutralization of a spike L452R SARS-CoV-2 variant. Cell 2021; 184: 3426–3437.e8.

48. Li Y, Ma M-L, Lei Q, Wang F, Hong W, Lai D-Y, et al. Linear epitope landscape of the SARS-CoV-2 Spike protein constructed from 1,051 COVID-19 patients. Cell Rep 2021; 34: 108915.

49. Singh A, Steinkellner G, Köchl K, Gruber K, Gruber CC. Serine 477 plays a crucial role in the interaction of the SARS-CoV-2 spike protein with the human receptor ACE2.

50. Hoffmann M, Kleine-Weber H, Pöhlmann S. A Multibasic Cleavage Site in the Spike Protein of SARS-CoV-2 Is Essential for Infection of Human Lung Cells. Mol Cell 2020; 78: 779–784.e5.

51. Papa G, Mallery DL, Albecka A, Welch L, Cattin-Ortolá J, Luptak J, et al. Furin cleavage of SARS-CoV-2 Spike promotes but is not essential for infection and cell-cell fusion.

52. Arora P, Sidarovich A, Krüger N, Kempf A, Nehlmeier I, Graichen L, et al. B.1.617.2 enters and fuses lung cells with increased efficiency and evades antibodies induced by infection and vaccination. Cell Rep 2021; 37: 109825.

53. Dutta NK, Mazumdar K, Gordy JT. The Nucleocapsid Protein of SARS–CoV-2: a Target for Vaccine Development. Journal of Virology . 2020., 94

54. Tylor S, Andonov A, Cutts T, Cao J, Grudesky E, Van Domselaar G, et al. The SR-rich motif in SARS-CoV nucleocapsid protein is important for virus replication. Canadian Journal of Microbiology . 2009., 55: 254–260

55. Leary S, Gaudieri S, Parker MD, Chopra A, James I, Pakala S, et al. Generation of a Novel SARS-CoV-2 Sub-genomic RNA Due to the R203K/G204R Variant in Nucleocapsid: Homologous Recombination has Potential to Change SARS-CoV-2 at Both Protein and RNA Level. Pathog Immun 2021; 6: 27–49.

56. Verba K, Gupta M, Azumaya C, Moritz M, Pourmal S, Diallo A, et al. CryoEM and AI reveal a structure of SARS-CoV-2 Nsp2, a multifunctional protein involved in key host processes. Res Sq 2021.

57. Rodrigues CHM, Pires DEV, Ascher DB. DynaMut2: Assessing changes in stability and flexibility upon single and multiple point missense mutations. Protein Sci 2021; 30: 60–69.

58. Jumper J, Tunyasuvunakool K, Kohli P, Hassabis D, AlphaFold Team. Computational predictions of protein structures associated with COVID-19. Deep Mind. https://deepmind.com/research/open-source/computational-predictions-of-protein-structures-associated-with-COVID-19.

59. Chen J, Malone B, Llewellyn E, Grasso M, Shelton PMM, Olinares PDB, et al. Structural Basis for Helicase-Polymerase Coupling in the SARS-CoV-2 Replication-Transcription Complex. Cell 2020; 182: 1560–1573.e13.

60. Gupta R, Charron J, Stenger CL, Painter J, Steward H, Cook TW, et al. SARS-CoV-2 (COVID-19) structural and evolutionary dynamicome: Insights into functional evolution and human genomics. J Biol Chem 2020; 295: 11742–11753.

61. Santerre M, Arjona SP, Allen CN, Shcherbik N, Sawaya BE. Why do SARS-CoV-2 NSPs rush to the ER? J Neurol 2020.

62. Thorne LG, Bouhaddou M, Reuschl A-K, Zuliani-Alvarez L, Polacco B, Pelin A, et al. Evolution of enhanced innate immune evasion by the SARS-CoV-2 B.1.1.7 UK variant. bioRxiv 2021.

63. Calistri P, Amato L, Puglia I, Cito F, Di Giuseppe A, Danzetta ML, et al. Infection sustained by lineage B.1.1.7 of SARS-CoV-2 is characterised by longer persistence and higher viral RNA loads in nasopharyngeal swabs. Int J Infect Dis 2021; 105: 753–755.

64. Yan L, Zhang Y, Ge J, Zheng L, Gao Y, Wang T, et al. Architecture of a SARS-CoV-2 mini replication and transcription complex. Nat Commun 2020; 11: 5874.

65. Jia Z, Yan L, Ren Z, Wu L, Wang J, Guo J, et al. Delicate structural coordination of the Severe Acute Respiratory Syndrome coronavirus Nsp13 upon ATP hydrolysis. Nucleic Acids Res 2019; 47: 6538–6550.

66. Jang K-J, Jeong S, Kang DY, Sp N, Yang YM, Kim D-E. A high ATP concentration enhances the cooperative translocation of the SARS coronavirus helicase nsP13 in the unwinding of duplex RNA. Sci Rep 2020; 10: 4481.

67. Guo G, Gao M, Gao X, Zhu B, Huang J, Luo K, et al. SARS-CoV-2 non-structural protein 13 (nsp13) hijacks host deubiquitinase USP13 and counteracts host antiviral immune response. Signal Transduct Target Ther 2021; 6: 119.

68. Zhang L, Jackson CB, Mou H, Ojha A, Peng H, Quinlan BD, et al. SARS-CoV-2 spike-protein D614G mutation increases virion spike density and infectivity. Nat Commun 2020; 11: 6013.

69. Korber B, Fischer WM, Gnanakaran S, Yoon H, Theiler J, Abfalterer W, et al. Tracking Changes in SARS-CoV-2 Spike: Evidence that D614G Increases Infectivity of the COVID-19 Virus. Cell 2020; 182: 812–827.e19.

70. Xia H, Cao Z, Xie X, Zhang X, Chen JY-C, Wang H, et al. Evasion of Type I Interferon by SARS-CoV-2. Cell Rep 2020; 33: 108234.

71. Garvin MR, T Prates E, Pavicic M, Jones P, Amos BK, Geiger A, et al. Potentially adaptive SARS-CoV-2 mutations discovered with novel spatiotemporal and explainable AI models. Genome Biol 2020; 21: 304.

72. Cedro-Tanda A, Gómez-Romero L, Alcaraz N, de Anda-Jauregui G, Peñaloza F, Moreno B, et al. The evolutionary landscape of SARS-CoV-2 variant B.1.1.519 and its clinical impact in Mexico City. bioRxiv. 2021.

73. Prates E, Garvin M, Jones P, Miller JI, Kyle S, Cliff A, et al. Antiviral Strategies Against SARS-CoV-2 – For a Bioinformatics Approach. In: Hann JJ, Bintou A, Keng C (eds). SARS-CoV-2 Methods and Protocols. 2021. Spriinger.

74. Katoh K, Misawa K, Kuma K-I, Miyata T. MAFFT: a novel method for rapid multiple sequence alignment based on fast Fourier transform. Nucleic Acids Res 2002; 30: 3059–3066.

75. Leigh JW, Bryant D. popart: full-feature software for haplotype network construction. Methods Ecol Evol 2015; 6: 1110–1116.

76. Shannon P, Markiel A, Ozier O, Baliga NS, Wang JT, Ramage D, et al. Cytoscape: a software environment for integrated models of biomolecular interaction networks. Genome Res 2003; 13: 2498–2504.

77. Ronquist F, Huelsenbeck JP. MrBayes 3: Bayesian phylogenetic inference under mixed models. Bioinformatics 2003; 19: 1572–1574.

78. Humphrey W, Dalke A, Schulten K. VMD: visual molecular dynamics. J Mol Graph 1996; 14: 33–8, 27–8.

79. The GIMP Development Team. (2019). GIMP. Retrieved from https://www.gimp.org. https://www.gimp.org.

80. Lindahl, Abraham, Hess, Spoel V der. GROMACS 2020 Source code. 2020.

81. Huang J, MacKerell AD Jr. CHARMM36 all-atom additive protein force field: validation based on comparison to NMR data. J Comput Chem 2013; 34: 2135–2145.

82. Guvench O, Mallajosyula SS, Raman EP, Hatcher E, Vanommeslaeghe K, Foster TJ, et al. CHARMM additive all-atom force field for carbohydrate derivatives and its utility in polysaccharide and carbohydrate-protein modeling. J Chem Theory Comput 2011; 7: 3162– 3180.

83. Huang P-S, Ban Y-EA, Richter F, Andre I, Vernon R, Schief WR, et al. RosettaRemodel: a generalized framework for flexible backbone protein design. PLoS One 2011; 6: e24109.

84. Jo S, Kim T, Iyer VG, Im W. CHARMM-GUI: a web-based graphical user interface for CHARMM. J Comput Chem 2008; 29: 1859–1865.

85. Jorgensen WL, Chandrasekhar J, Madura JD, Impey RW, Klein ML. Comparison of simple potential functions for simulating liquid water. J Chem Phys 1983; 79: 926–935.

86. Evans DJ, Holian BL. The Nose–Hoover thermostat. J Chem Phys 1985; 83: 4069–4074.

87. Parrinello M, Rahman A. Polymorphic transitions in single crystals: A new molecular dynamics method. J Appl Phys 1981; 52: 7182–7190.

88. Darden T, York D, Pedersen L. Particle mesh Ewald: An N⋅log(N) method for Ewald sums in large systems. J Chem Phys 1993; 98: 10089–10092.

89. Hess B, Bekker H, Berendsen HJC, Fraaije JGEM. LINCS: A linear constraint solver for molecular simulations. J Comput Chem 1997; 18: 1463–1472.

