## Supplementary Tables for "Rapid expansion of SARS-CoV-2 variants of concern is a result of adaptive epistasis"

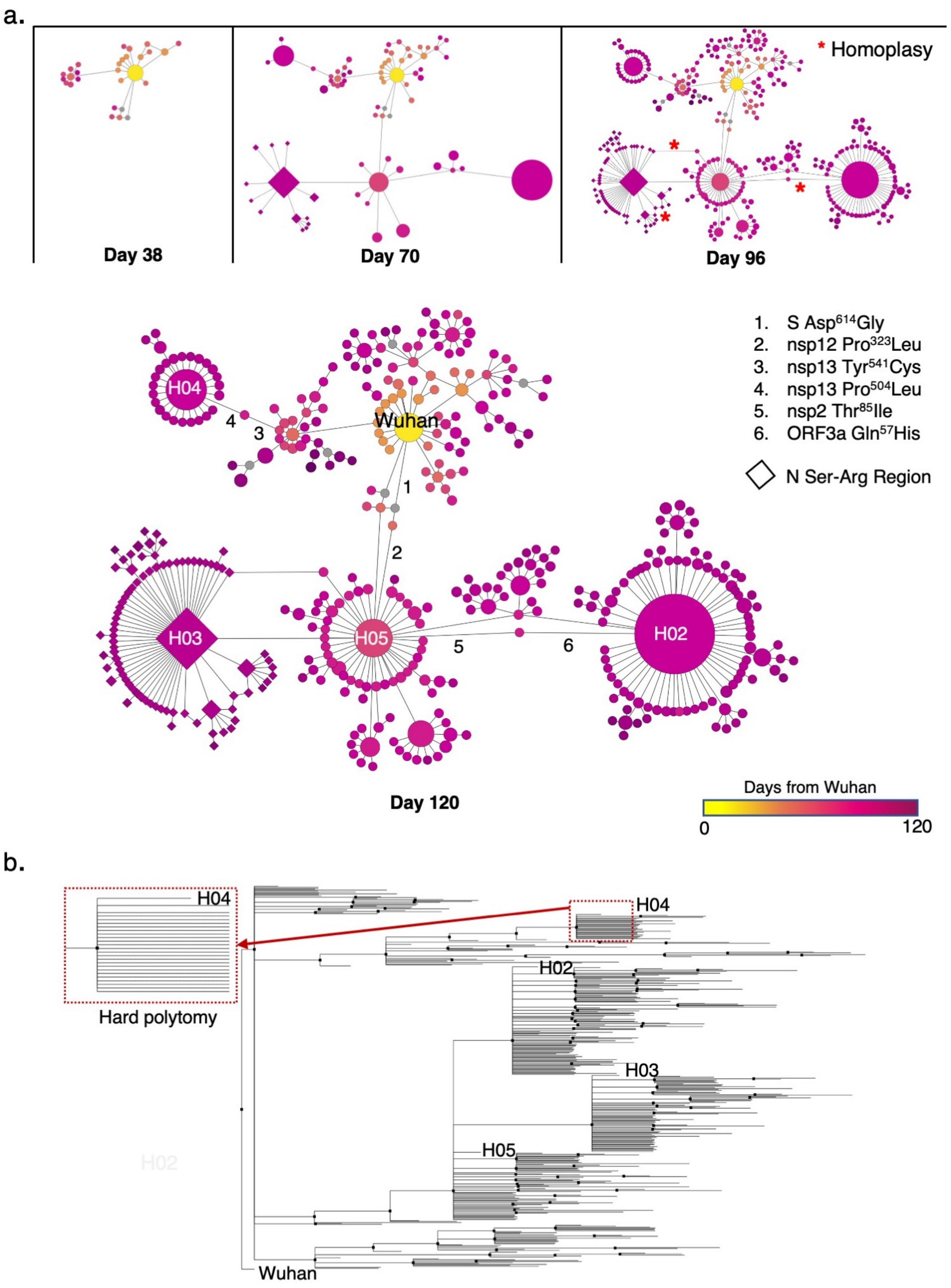
a.
* Homoplasy
Day 38
Day 70
Day 96
1. S Asp$^{614}$Gly
2. nsp12 Pro$^{323}$Leu
3. nsp13 Tyr$^{541}$Cys
4. nsp13 Pro$^{504}$Leu
5. nsp2 Thr$^{85}$Ile
6. ORF3a Gln$^{57}$His
N Ser-Arg Region
Wuhan
H04
H03
H05
H02
1
2
3
4
5
6
Day 120
Days from Wuhan
0
120
b.
H04
Hard polytomy
H04
H02
H03
H05
Wuhan

*Probability of mutation accumulation*

To calculate the chance of accumulating several mutations in a certain period, the probability density function for a normal distribution is used:

$$PDF(x) = exp(-(x-\mu)^2/2\sigma^2)/sqrt(2\pi * \sigma^2),$$

**Supplementary Figures**

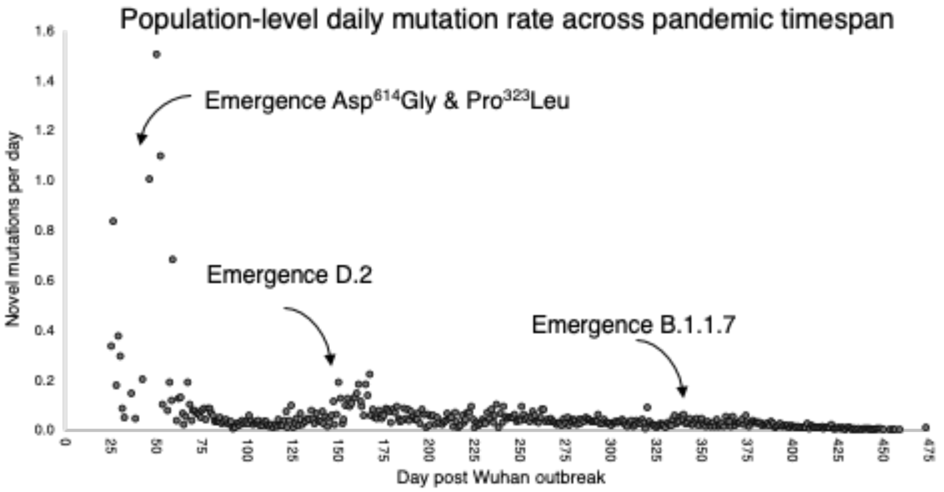


**Fig. S1.**

**Population level mutation rate over the course of the pandemic.** Number of novel mutations sampled across the globe for each day are plotted against time (days from the Wuhan outbreak). Emergence of major VOC are provided for context and show small increases in the number of new mutations but there is an overall decrease across time, even accounting for multiple mutations at a site to different nucleotide states and deletions.

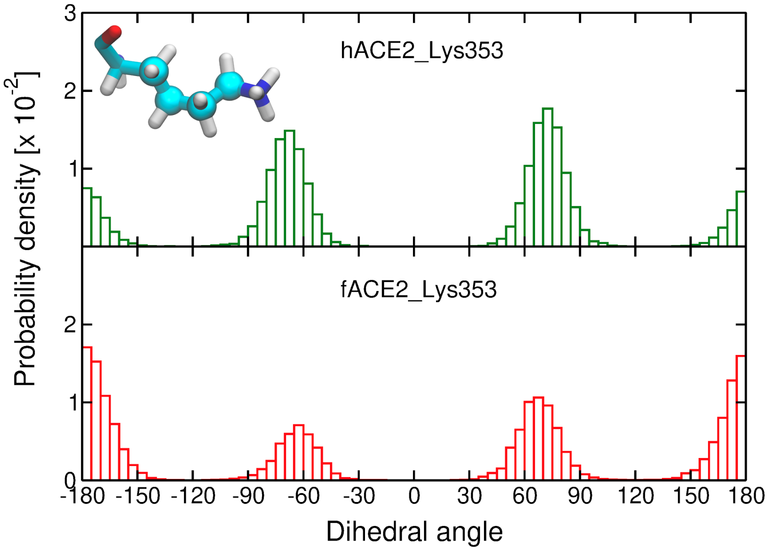


**Fig. S2.**

**The probability density of the conformations of Lys$^{353}$ in human and ferret ACE2 in the simulations.**
Histograms of the distribution of a dihedral angle of the Lys$^{353}$ side chain carbon atoms in human ACE2 (hACE2, upper figure) and ferret ACE2 (fACE2, lower figure) in complex with the SARS-CoV-2 S receptor-binding domain. The atoms forming the selected dihedral are depicted as spheres in the molecular representation of Lys$^{353}$. Three independent simulations are considered for the calculation of the histograms. Dihedral angles near ±180$^{o}$ correspond to a more stretched conformation (i.e., *trans*).

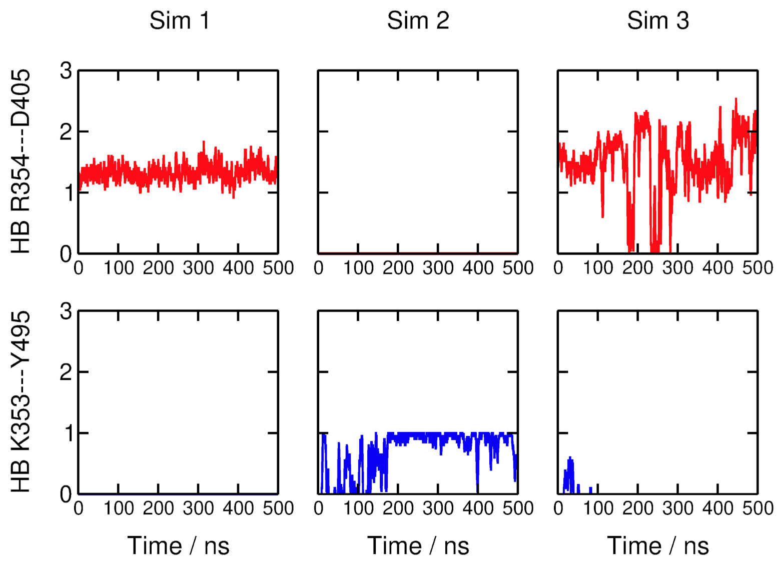


**Fig. S3.**

**Competing hydrogen bond interactions formed between positively charged amino acid residues in ferret ACE2 (fACE2) and the SARS-CoV-2 S receptor-binding domain.** Time evolution of the number of hydrogen bonds (HB) that fACE2 Arg3$^{54}$ and Lys$^{353}$ form with Asp$^{405}$ and Tyr$^{495}$ from the SARS-CoV-2 S receptor-binding domain. The columns correspond to the three simulation replicas. The geometric criteria adopted for hydrogen bonds are a cutoff of 3.0 Å for donor-acceptor distance and 20º for acceptor-donor-H angle.

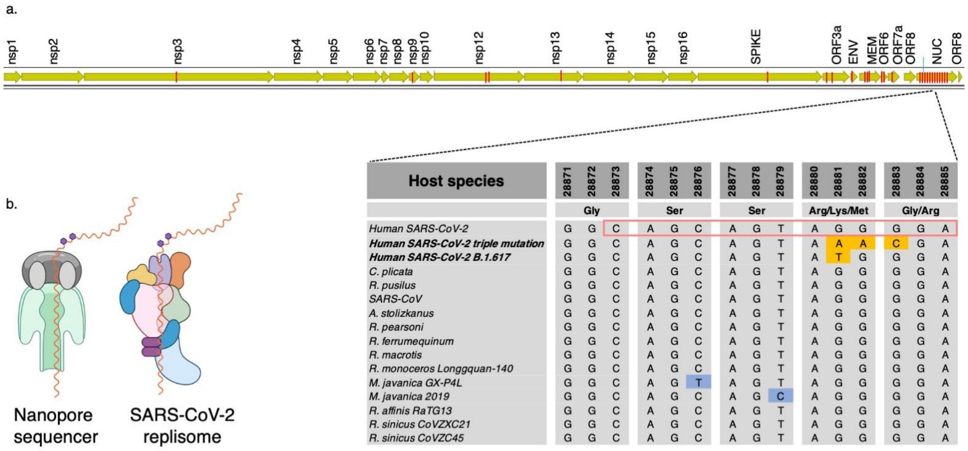


**Fig. S4**

**Modifications at the Ser-Arg-rich region of N may affect replication speed. a.** Location of 41 epigenetic sites reported in Kim et al. 2020 (red bars on SARS-CoV-2 genome). One of the sites in the nucleocapsid gene (nucleotides in red box of aligned sequences) is highly conserved across diverse host-defined coronaviruses. All bats and human coronavirus species from China are completely conserved at the epigenetic site 28881-28883, except for a 3-bp mutation in SARS-CoV-2 that occurred early in the pandemic and now corresponds to ~50% of all sequences globally (diamond nodes in Figure 1). **b.** Kim et al. proposed that $N^6$-methyladeonsine modification of the genome (purple hexagons), common in RNA viruses, caused the strand to pause while traversing the nanopore sequencing apparatus. We propose that loss of this site via mutations at site 203 in N may increase the replication rate of the RNA strand through the SARS-CoV-2 replisome. *Aselliscus stoliczkanus* - Stoliczka's trident bat, *Chaerephon plicata* - wrinkle-lipped free-tailed bat, *Rhinolophus pusillus* - least horseshoe bat, *R. pearsoni* - Pearson's horseshoe bat, *R. macrotis* - big-eared horseshoe bat, *R. ferrumequinum* - greater horseshoe bat, *R. monoceros* - Formosan lesser horseshoe bat, *R. affinis* intermediate horseshoe bat, *R. sinicus* Chinese rufous horseshoe bat, *R. mayalanis* - Mayalan horseshoe bat, *SARS* - Severe Acute Respiratory Syndrome, *Manis javanica* - Malayan pangolin. *Created with BioRender.com.*
